# Systematic contextual biases in SegmentNT potentially relevant to other nucleotide transformer models

**DOI:** 10.1101/2025.04.09.647946

**Authors:** Mark T. W. Ebbert, Anna Ho, Madeline L. Page, Bram Dutch, Blake K. Byer, Kristen L. Hankins, Hady Sabra, Bernardo Aguzzoli Heberle, Mark E. Wadsworth, Grant A. Fox, Bikram Karki, Caylin Hickey, David W. Fardo, Cody Bumgardner, Yasminka A. Jakubek, Cody J. Steely, Justin B. Miller

**Author notes:** To whom correspondence should be addressed: Mark Ebbert, Justin Miller. Author Disclosures: None. These authors contributed equally and share co-first authorship (Mark T. W. Ebbert, Anna Ho, Madeline L. Page).

## Abstract

Recent advances in large language models (LLMs) have extended to genomic applications, yet model robustness relative to context is unclear. Here, we demonstrate two intrinsic biases (input sequence length and nucleotide position) affecting SegmentNT results, a model included with the Nucleotide Transformer that provides nucleotide-level predictions of biological features. We demonstrate that nucleotide position within the input sequence (beginning, middle, or end) alters the nature of SegmentNT’s raw prediction probabilities, which can be standardized to improve prediction consistency. While longer input sequence length improves model performance, diminishing returns suggest a surprisingly small input length of ∼3,072 nucleotides might be sufficient for many applications. We further identify a 24-nucleotide periodic oscillation in SegmentNT’s prediction probabilities, revealing an intrinsic bias potentially linked to the model’s training tokenization (6-mers) and architecture. We identify potential approaches to account for these biases and provide generalizable insights for utilizing nucleotide-resolution functional prediction models.

---

Foundation models are powerful artificial intelligence models pretrained on large datasets to capture generalized knowledge that can be fine-tuned for specific tasks^1,2^. Such models have significantly advanced multiple fields, including natural language processing^3–6^, computer vision^7–10^, graph learning^11–14^, and protein folding prediction^15^, by leveraging architectures such as convolutional neural networks (CNNs)^16–24^, long short-term memory (LSTM) networks^25–28^, and transformers^29,30^. These successes have driven interest in developing foundation models tailored for genomic applications^31–39^.

One such model, SegmentNT^38^, is a fine-tuned version of the Nucleotide Transformer^39^, developed to predict 14 genomic features at single-nucleotide resolution. Unlike most genomic foundation models that provide sequence-wide predictions, SegmentNT refines its predictions down to individual nucleotides, enabling greater precision in functional annotation. However, little is known about potential underlying biases in that approach and how they affect model performance.

Here, to contribute to improving the utility of powerful models like SegmentNT, we describe two systematic biases—input sequence length and nucleotide position, therein—affecting SegmentNT performance using two of SegmentNT’s feature predictions (intronic and exonic probabilities). We only focus on these two feature predictions because: (1) they are of broad interest, (2) we wanted to provide more in-depth analyses, and (3) they were sufficient to demonstrate the biases. We demonstrate that nucleotide position within the input sequence significantly alters the nature of SegmentNT’s raw probabilities, introducing variability depending on whether the nucleotide appears at the beginning, middle, or end. As alluded to in the SegmentNT publication^38^, we also show that increasing sequence length improves model performance but with diminishing returns, suggesting an optimal input size much smaller than the maximum model input, depending on user needs. We further identify notable discrepancies between SegmentNT and gene annotations that may indicate the model is identifying additional biological characteristics, but further work is needed to determine whether these predictions identify new biology or are inaccuracies. Finally, we identify a 24-nucleotide periodic oscillation in SegmentNT’s probabilities, revealing an inherent bias potentially linked to the model’s training tokenization (6-mers), rotary positional embeddings (RoPE), and U-Net architecture.

SegmentNT and other models demonstrate the power of artificial intelligence and will serve as a major boon to genomic research. Understanding these biases is critical for end users to properly interpret results from SegmentNT (and potentially other genomic foundation models) and to improve training. Our findings provide actionable and generalizable insights to optimize sequence selection for downstream applications, improve foundation model training, and mitigate biases in these important and valuable genomic foundation models.

## Results

### Nucleotide position within input sequence significantly biases raw SegmentNT probabilities

We selected the five most-studied human genes^40^ (GRCh38) with high biological significance (*APOE, EGFR, TP53, TNF*, & *VEGFA*; **Fig. 1**) and a non-genic region (chr3:153050001-153053598; negative control sequence) to assess whether nucleotide position within the input sequence biases SegmentNT’s probabilities. SegmentNT predicts 14 attributes for individual nucleotides^38^, but we focused on two representative attributes: intronic and exonic probabilities. Those probabilities were calculated individually and were not constrained by the other predictions (i.e., the exon and intron predictions did not sum to 1). Specifically, we assessed whether the probabilities varied, depending on whether the nucleotide was the first, middle, or last nucleotide in the input sequence. Here, we used 24,576 nucleotide input sequences (4,096 tokens of six nucleotides).

**Fig 1.**
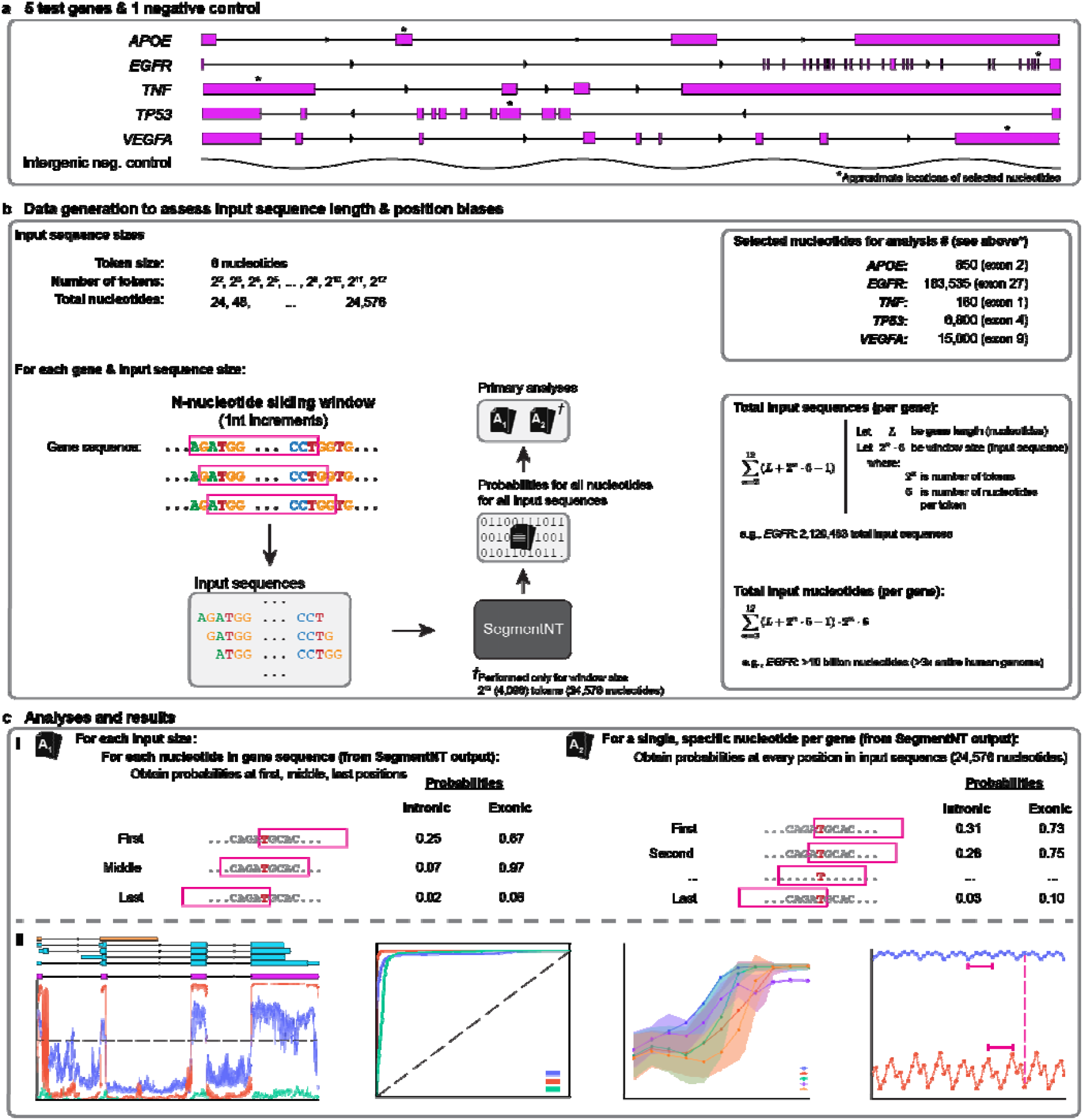
Graphical abstract illustrating how we assessed two intrinsic biases in SegmentNT. (**a**) We selected the five most-studied genes (*APOE, EGFR, TNF, TP53*, & *VEGFA*) with known biological significance along with a single intergenic region (chr3:153050001-153053598; negative control) as test regions to formally assess two intrinsic biases in SegmentNT: (1) input sequence length, and (2) nucleotide position, therein. For downstream analyses, we also arbitrarily selected a single nucleotide from each gene; approximate locations indicated by asterisks (*). *TP53* exons shown in reverse order (right to left) because the gene is on the negative strand. (**b**) Diagram illustrating how we generated SegmentNT probabilities across eleven window input sizes (1-nucleotide increments; nt). For each genomic region, we generated input sequences for each distinct sliding window sizes, incrementing the number of tokens (1 token = 6 nucleotides) by 2^n^, beginning with 2^2^ tokens (4 tokens * 6 nucleotides per token = 24 nucleotides) through 2^12^ (4,096 tokens; 24,576 nucleotides). Total number of input sequences per gene equaled the length of the gene (in nucleotides) + length of the window – 1, for all window sizes (e.g., *EGFR* for window size 24,576: 188,306 + 24,576 – 1 = 212,881 input sequences; total input sequences: 2,120,483; total input bases: 10,056,354,216). From SegmentNT results, we performed two primary analyses (indicated by **A**_**1**_ and **A**_**2**_; see figure **c**). Exact nucleotides selected for deeper analysis in **Analysis A**_**2**_ are provided. (**c**) Illustrations and representative figures of primary analyses (**A**_**1**_ and **A**_**2**_) and results. (**c**_**i**_) **Analysis A**_**1**_: for each gene and window input size we assessed nucleotide position bias based on every nucleotide’s probability when it was first, middle, or last within the input sequence. **Analysis A**_**2**_: we assessed nucleotide position bias across the *entire* input sequence for the five specific nucleotides indicated in **a** and **b**. (**c**_**ii**_) Representative figures of downstream interpretive analyses illustrating how both biases affect SegmentNT’s raw probabilities and model performance.

For *APOE*, which harbors the strongest known genetic risk factor for Alzheimer’s disease^41–43^, the middle position initially appeared most accurate when taking the probabilities at face value (**Fig. 2a**; see **Supplementary Data** for interactive plots). Specifically, nucleotides within canonical NCBI RefSeq^44^ exons had high exonic probabilities (median = 0.993) while nucleotides within canonical introns had low exonic probabilities (median = 0.023). Because SegmentNT’s exonic probabilities for nucleotides in the middle position were mostly dichotomous (i.e., near 1 for nucleotides within exons and near 0 for nucleotides within introns), we selected an intuitive threshold of 0.50 (halfway between 0 and 1) to estimate predictive accuracy, achieving 98.3% accuracy. Sensitivity, specificity, and Area Under the Curve (AUC) were 98.6%, 98.2%, and 0.9990, respectively (**Supplemental Fig. 1**).

**Figure 2.**
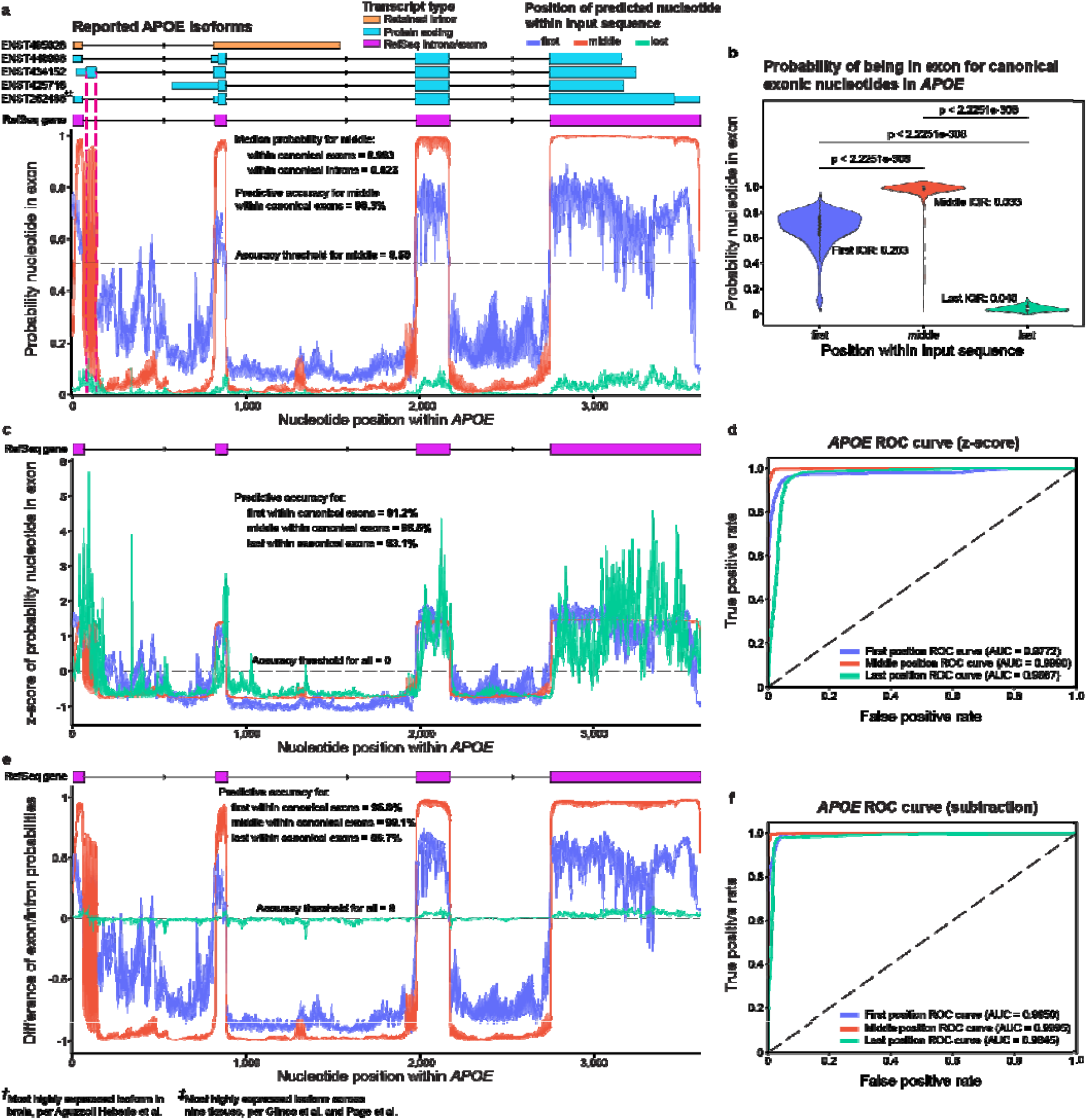
Nucleotide position within input sequence significantly biases SegmentNT probabilities in *APOE*. **(a)** Plot showing exonic probabilities for each *APOE* nucleotide. All three positions result in distinct distributions that require unique interpretation. At face value, middle position appeared most intuitive and accurate. Canonical RefSeq *APOE* exons are shown, as are all reported Ensembl isoforms, including which isoform is most highly expressed in the human frontal cortex^45^ and across nine GTEx tissues^46,47^; shorter boxes indicate untranslated regions (UTRs), while taller boxes identify protein-coding sequences. One region shows a strong but vacillating signal that aligns with an alternatively spliced exon in RNA isoform ENST00000434152 (Supplemental Fig 10). (**b**) Violin plots showing exonic probability distributions (first, middle, last) for exonic nucleotides (per RefSeq exons). Middle resulted in higher probabilities than first (p < 2.2251e-308) and last (p < 2.2251e-308). Last resulted in lower probabilities than first (p < 2.2251e-308). Exact p-values could not be calculated because the p-values were lower than the lowest reportable p-value in Python. First resulted in greater variability, based on interquartile range (IQR). (**c**) Plot showing Z-scores of exonic probabilities for each *APOE* nucleotide. All three positions provided high predictive value, but the middle position was most accurate (z = 0; 98.5%; first = 91.2%; last = 93.1%). (**d**) Receiver Operating Characteristic (ROC) curve based on Z-score showing True vs. False Positive rates for first, middle, and last positions, and their respective Areas Under the Curve (AUCs). (**e**) Plot showing normalized exonic probabilities (exon – intron) for each *APOE* nucleotide. (**f**) ROC curve based on subtraction model. RNA isoforms were plotted using RNApysoforms^48^.

Exonic probabilities for the first and last position appeared less reliable than middle at face value, as first resulted in lower and higher exonic probabilities within exons and introns, respectively, while last had consistently low exonic probabilities for all nucleotides (**Fig. 2a,b**). The middle position had significantly higher exonic probabilities within exons compared to when the same nucleotide was in the first (p < 2.23e-308) or last (p < 2.23e-308; **Fig. 2b**) position; exact p-values could not be provided because they were lower than the current lowest reportable p-value in Python, which is approximately 2.23e-308 based on Python’s minimum positive float value (`sys.float_info.min`). Based on interquartile range (IQR), variability of the probabilities was greater for first (IQR = 0.203) than middle (IQR = 0.033) or last (IQR = 0.040). For intronic nucleotides, middle exonic probabilities were lower than first (p = 1.65e-296) and higher than last (p = 3.68e-79) positions (**Supplemental Fig. 2**).

Although raw probabilities differed significantly across positions, upon deeper investigation, all three provided high predictive value, demonstrating that the model is inherently robust but requires additional standardization for uniform interpretation. To standardize interpretation, we normalized probabilities across input sequence positions and assessed model performance for first, middle, and last positions. As proof-of-principle, we tested two simplistic normalization methods—Z-score (**Fig.2c**) and subtraction (exonic minus intronic probability, **Fig. 2e**)—both of which effectively standardized interpretation.

Z-score ultimately puts all three score sets on the same scale, resulting in nearly identical performance metrics as raw scores, but can be interpreted uniformly across positions. Thus, the first and last exhibited greater variability of the probabilities (**Fig. 2c**). Middle remained most accurate (98.5%; **Fig. 2c**), using Z = 0 to assess accuracy across all three positions (**Fig. 2c**), and had the greatest sensitivity (99.1%) and specificity (98.2%) for exonic probabilities of exonic nucleotides (AUC = 0.9990; **Fig. 2d**) compared to first (accuracy = 91.2%; sensitivity = 97.0%; specificity = 88.3%; AUC = 0.9772) and last positions (accuracy = 93.1%; sensitivity = 90.0%; specificity = 94.6%; AUC = 0.9667).

The subtraction model performed similarly. Middle had the greatest accuracy (99.1%; **Fig. 2e**), sensitivity (99.8%), and specificity (98.8%; AUC = 0.9995; **Fig. 2f**) compared to first (accuracy = 96.8%; sensitivity = 98.0%; specificity = 96.2%; AUC = 0.9950) and last (accuracy = 96.7%; sensitivity = 98.1%; specificity = 96.0%; AUC = 0.9845), using the normalized threshold of 0 to assess accuracy.

Results were similar when predicting whether a given nucleotide was intronic (**Supplemental Fig. 3a**). Intronic probabilities between the middle position and the first (p < 1.58e-296) and last (p < 2.23e-308) positions were significantly different (**Supplemental Fig. 3b**).

The other four genes (**Fig. 3a-d**) and the non-genic region (**Supplemental Fig. 4**) also exhibited clear differences in probabilities, depending on whether the nucleotide was first, middle, or last in the input sequence. As validation, we tested five additional gene bodies (*DPM2, ECM1, LINC00207, NAV2-AS5, WFDC5*), two of which are non-coding RNAs (*LINC00207 & NAV2-AS5*); all five also exhibited clearly distinct distributions for first, middle, and last positions (**Supplemental Figs. 5-9**). Thus, although all three positions provide high predictive value, these data clearly demonstrate that a nucleotide’s location within the input sequence significantly biases SegmentNT’s raw probabilities and that the nature of the probability distributions is distinct across positions. Thus, interpreting and leveraging SegmentNT’s nucleotide-resolution in future applications must account for this intrinsic bias. Our results further suggest, however, that the underlying model is surprisingly robust since even nucleotides at the beginning and end of the input sequence (i.e., lacking valuable up or downstream sequence context) still provide high predictive value when properly interpreted. These results may also be applicable to future nucleotide foundation models utilizing a similar architecture as the Nucleotide Transformer and SegmentNT.

**Fig 3.**
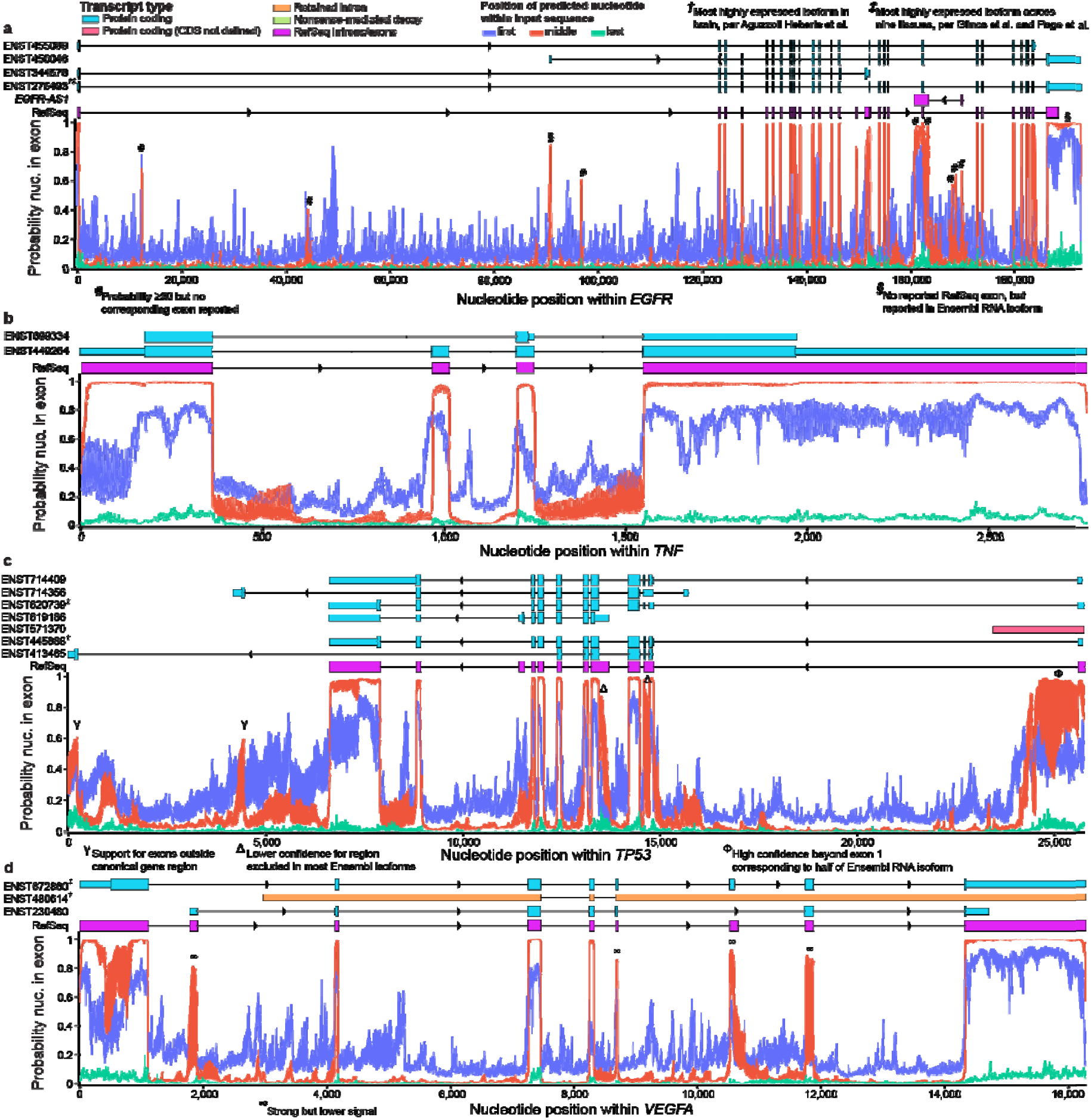
Nucleotide position significantly biases SegmentNT probabilities in four additional genes. To assess whether the results from Figure 1 are likely to generalize across the genome, we assessed nucleotide position in four additional genes (*EGFR, TNF, TP53*, & *VEGFA*), and a negative control sequence (chr3:153050001-153053598; Supplemental Fig. 4). (**a**) Plot showing SegmentNT exonic probabilities for each *EGFR* nucleotide. Four representative *EGFR* RNA isoforms (13 total) and the canonical RefSeq exons are shown. Several discrepancies between SegmentNT’s predictions and either RefSeq or Ensembl RNA isoforms are indicated; some correspond to *EGFR-AS1* exons on the negative strand. Thus, SegmentNT results should not be interpreted through the perspective of a single gene or strand. (**b**) SegmentNT exonic probabilities for *TNF*, along with both reported RNA isoforms and RefSeq exons. Neither isoform reached our minimal expression threshold (median counts-per-million ≥ 1) to report one as being most expressed in any tissue, likely because *TNF* is expressed as a response to inflammation and cellular stress. (**c**) SegmentNT exonic probabilities for *TP53*. Seven representative reported Ensembl RNA isoforms (33 total) and RefSeq exons are also shown. Four *TP53* isoforms extend beyond the canonical RefSeq gene boundaries (two shown; see Supplemental Fig. 14). *TP53* exons are in reverse order because it is on the negative strand of the human GRCh38 reference genome. (**d**) SegmentNT exonic probabilities for *VEGFA*, along with three representative RNA isoforms (26 total) and RefSeq exons. ENST00000672860 was most expressed across the nine GTEx tissues used in Page et al., per data from Glinos et al.^46,47^ Surprisingly, an isoform with a reported retained intron (ENST00000480614) was most expressed in human frontal cortex, per Aguzzoli Heberle et al.^45^ There were several *VEGFA* exons with lower, albeit strong, probabilities from SegmentNT. Why these exons have lower probabilities is unclear. For all genes, we selected at least one representative isoform for each reported exon. For isoforms, shorter boxes identify untranslated regions (UTRs), while taller boxes identify protein-coding sequence. RNA isoforms were plotted using RNApysoforms^48^.

### SegmentNT exonic probabilities may capture additional biologically meaningful information

One of the potential promises of artificial intelligence is to help identify patterns and associations previously overlooked by traditional methods (i.e., hypothesis generation). Thus, we assessed perceived discrepancies between SegmentNT’s exonic probabilities and RefSeq’s canonical exon annotations; as previously demonstrated^38^, SegmentNT’s exonic probabilities generally concurred with annotations, but we observed several deviations that may suggest SegmentNT is capturing additional biological information. Whether these discrepancies are inaccurate predictions or inaccurate gene annotations remains unclear, but regardless, they may be some of the most important results to be studied because they either represent a collection of inaccurate results that could improve future models, or they represent additional biological information gained learned through deep model training (i.e., new hypotheses). For example, several groups recently discovered hundreds to thousands of new RNA isoforms using long-read sequencing^45,47,49^, demonstrating many genomic mysteries remain.

Within *APOE*, SegmentNT indicated a strong but inconsistent exonic signal between approximately nucleotides 61 and 143, aligning with an extension of exon 1, as demonstrated by Ensembl^50^ *APOE* isoform ENST00000434152, which extends from nucleotide 61 to 133 (**Fig. 2a**; **Supplemental Fig. 10**). SegmentNT’s exonic probabilities for this region oscillate dramatically within four nucleotides, ranging from 0.06 to 0.96; why it oscillates so dramatically is unclear, but it could indicate alternative splicing or an uncharacterized functional region. For interest, the most expressed RNA isoform in human frontal cortex, per Aguzzoli Heberle et al.^45^, was ENST00000252486 (**Fig. 2a; Supplemental Fig. 11**), which aligns to the canonical RefSeq exons; this isoform was also the most expressed across nine GTEx tissues per Glinos et al.^47^, based on analyses by Page et al.^46^ (**Supplemental Fig. 11**).

In *EGFR*, we observed several spikes (exonic probabilities >0.3; clearly above noise signal) that did not match reported RefSeq or Ensembl *EGFR* exons (indicated by # in **Fig. 3a**; all 13 reported isoforms shown in **Supplemental Fig. 12**); some aligned with *EGFR-AS1* on the opposite strand, highlighting that SegmentNT predictions should not be interpreted through the perspective of a single gene or strand. Two spikes corresponded to RNA isoform exons not reported by RefSeq (§) that have since been added to RefSeq version 40, including the last exon being extended (**Fig. 3a**). In both data sets, the most expressed *EGFR* RNA isoform was ENST00000275493^45–47^ (**Supplemental Fig. 13**). SegmentNT concurred with canonical *TNF* exons (**Fig. 3b**).

For *TP53*, four reported Ensembl RNA isoforms extend well beyond RefSeq gene boundaries. Two exons outside canonical *TP53* gene boundaries were moderately supported (>0.50; **Fig. 3c;** Y), while the others (annotated as “nonsense-mediated decay”) lacked support (**Supplemental Fig. 14**). SegmentNT may have also indicated alternative splicing, with strong but lower probabilities where exon usage varies across isoforms (e.g., RefSeq exons 2 and 4 from right to left; Δ). For example, RefSeq exon 2 is only sometimes spliced into two exons. Similarly, SegmentNT indicates higher probabilities between RefSeq exons 1 and 2 (Φ) that correspond to isoform ENST00000571370 annotated as “protein coding (CDS not defined)”, though SegmentNT did not support its full length. The most expressed isoforms in the frontal cortex were ENST00000445888 and ENST00000619485, while ENST00000620739 was most expressed across GTEx tissues (**Supplemental Fig. 15**).

In *VEGFA*, SegmentNT indicated lower probabilities for several exons (**Fig. 3d**; ∞; **Supplemental Fig. 16**). Surprisingly, the most expressed *VEGFA* RNA isoform in human frontal cortex was ENST00000480614^45^ (**Supplemental Fig. 17**), which has a retained intron. These discrepancies provide useful case studies for future study. As expected, exonic probabilities within the non-genic region did not indicate any exons within the region (all probabilities < 0.3; **Supplemental Fig. 4**).

### Longer input sequence improves model performance, with diminishing returns

After assessing the nucleotide position bias, we assessed input sequence length. As expected, and as shown in the original SegmentNT paper^38^, longer input sequences resulted in higher probabilities, though with diminishing returns. Identifying a lower threshold that provides sufficient performance for a given task while minimizing computational time may be important in resource-limited situations. We assessed eleven input lengths ranging from 24 to 24,576 nucleotides for exonic *APOE* nucleotides (**Fig. 4a-e**; **Supplemental Table 1**). SegmentNT’s base model (Nucleotide Transformer^39^) was trained on six-nucleotide tokens, requiring input sequences to be multiples of six. We assessed tokens in increments of 2^n^, beginning with 2^2^ (4, 8, 16, …, 4096).

**Fig. 4.**
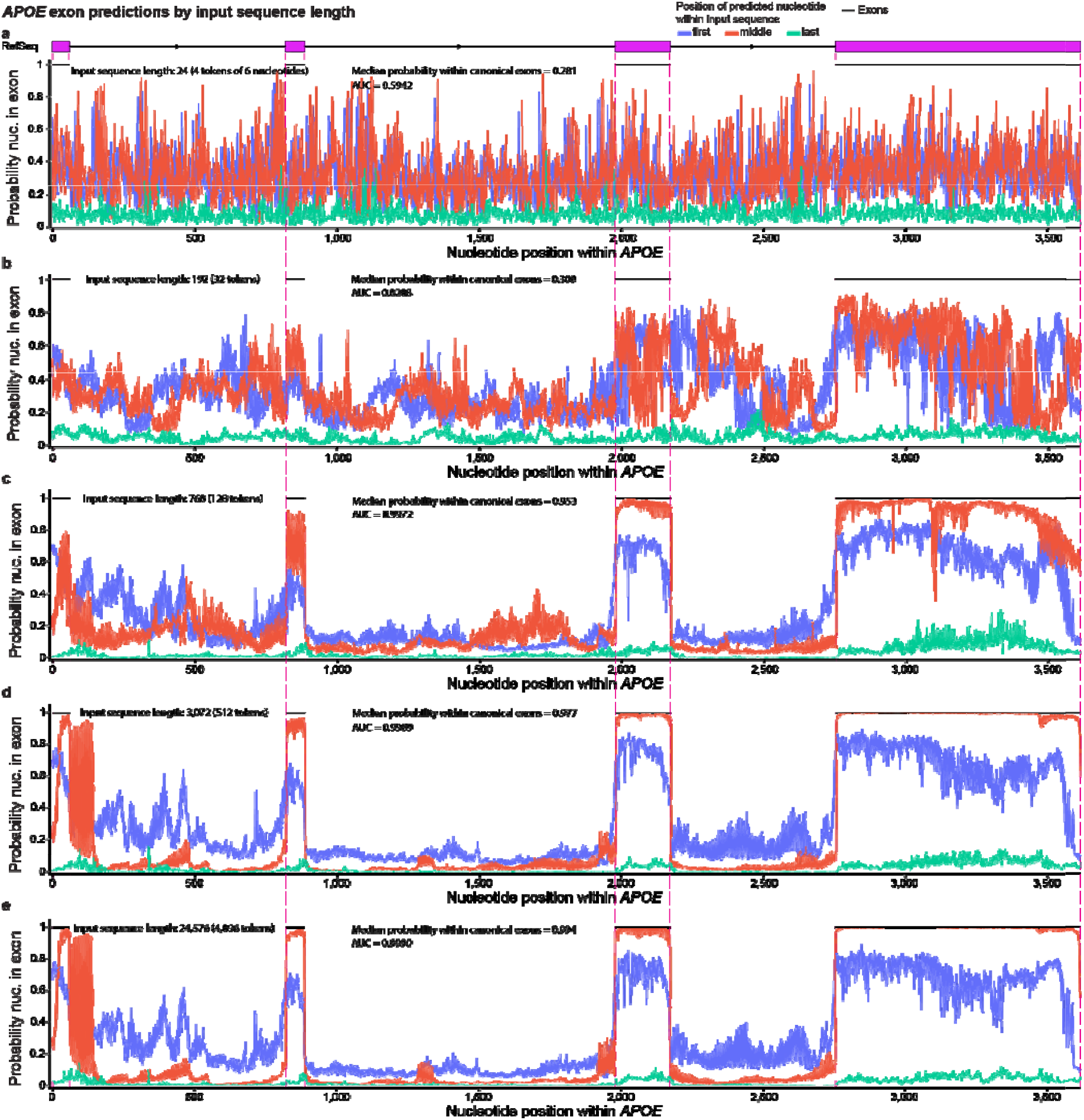
Longer input sequence increases model performance, with diminishing returns. We assessed eleven input lengths ranging from 24 to 24,576 nucleotides and found that longer input improves performance, based on median exonic probabilities and Areas Under the Curve (AUCs). SegmentNT’s base model (Nucleotide Transformer) was trained on six-nucleotide tokens, requiring input in multiples of six (one token). SegmentNT requires the number of tokens within the input sequence to be in increments of 2^n^ (4, 8, 16, …, 4096). Results for five representative input lengths are shown. Median exonic probabilities and model AUCs for the middle position are reported. (**a**) SegmentNT exonic probabilities for each *APOE* nucleotide using a 24-nucleotide input sequence (4 tokens). No distinguishable pattern between intronic and exonic nucleotides exists. (**b**) Same as figure **a**, but with a 192-nucleotide input sequences (32 tokens). Probabilities appear less random, but without a clear distinction between intronic and exonic nucleotides. (**c**) 768-nucleotide input sequence (128 tokens). A clearly non-random pattern distinguishing intronic and exonic nucleotides emerges as the model AUC jumps to 0.9972. (**d**) 3,072-nucleotide Input sequence (512 tokens). Probabilities further stabilize. A strong but vacillating signal is clear just beyond the first exon, which corresponds to an alternative exon found in a reported Ensembl RNA isoform (see Figure 1, Supplemental Figure 10). (**e**) 24,576-nucleotide input sequence (4096 tokens). Exonic probabilities and model AUC approach one, while probabilities for nucleotides within canonical RefSeq introns approach zero.

Exonic probabilities for 24-nucleotide sequences showed no distinguishable pattern between exons and introns (**Fig. 4a**). With 192 nucleotides, the pattern remains indistinguishable, but probabilities appear less sporadic (**Fig. 4b**). With 768 nucleotides, a clearly non-random pattern emerges between exons and introns (**Fig. 4c**). The respective probabilities and AUCs increased with sequence length, but with diminishing returns (**Fig. 4d,e**; **Fig. 5**; **Supplemental Tables 1**,**2**). Using AUC, the most benefit was generally achieved by an input sequence of 3,072 nucleotides for exonic predictions, though larger studies will be necessary to reliably identify the ideal input threshold to minimize computational costs (**Fig. 5b**; **Supplemental Table 2**). Although the original SegmentNT paper^38^ found additional benefit with longer input sequences, the benefit diminishes in their results, also. We observed similar trends for the other four primary genes tested (**Supplemental Figs. 18-21**; **Fig. 5**; **Supplemental Tables 1**,**2**), demonstrating that: (1) input length matters, and (2) longer input sequences perform better, but with minimal gains after ∼3,072 nucleotides, based on AUC using the raw predictive values.

**Fig 5.**
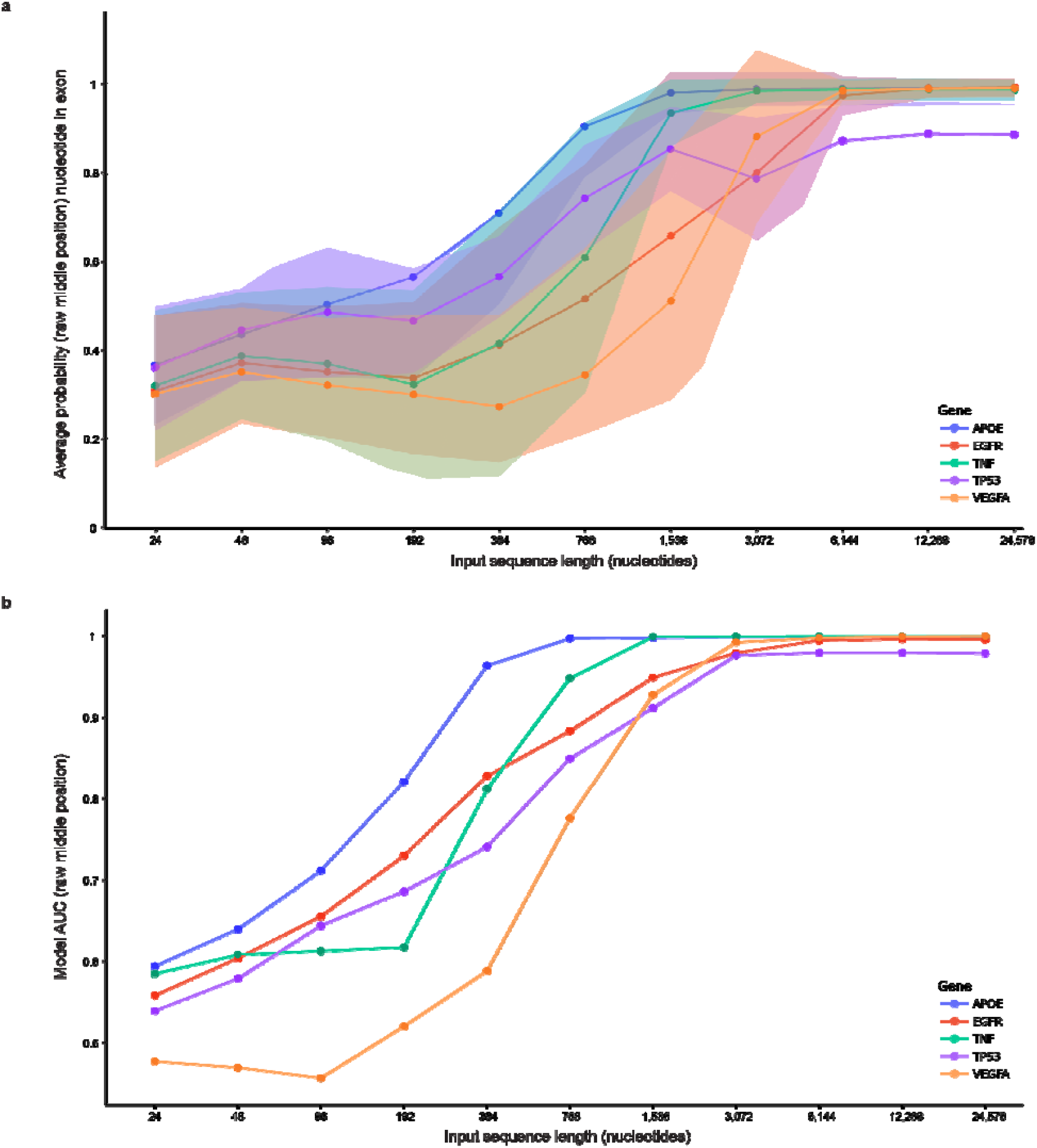
Smaller gains in model performance achieved beyond an input sequence length of 3,072 nucleotides. (**a**) Based on raw mean probabilities (middle position), the most benefit in maximizing SegmentNT probabilities is achieved by input sequence length of 6,144 nucleotides, as this is where the increase plateaus. Standard deviations are plotted for each gene. (**b**) Based on the Areas Under the Curves (AUCs) using the raw probabilities for middle position, most gain in model performance is achieved by length 3,072 nucleotides for all five genes. For *VEGFA*, model performance is <0.5 for input lengths ≤96 nucleotides. These results suggest that most benefit is achieved with an input sequence of 3,072 nucleotides since AUC provides a more meaningful measure of model performance than mean probabilities.

### SegmentNT probabilities oscillate in 24-nucleotide intervals

We next formally assessed the stability of probabilities across the full window of a 24,576-nucleotide input sequence. We quantified SegmentNT’s probabilities for a single nucleotide at every position within the input sequence (24,576 times) using *APOE* nucleotide 850 (exon 2; **Fig. 6a**). For example, for the first input sequence, nucleotide 850 was the first nucleotide in the input; for the tenth input, nucleotide 850 was tenth, etc.

**Fig. 6.**
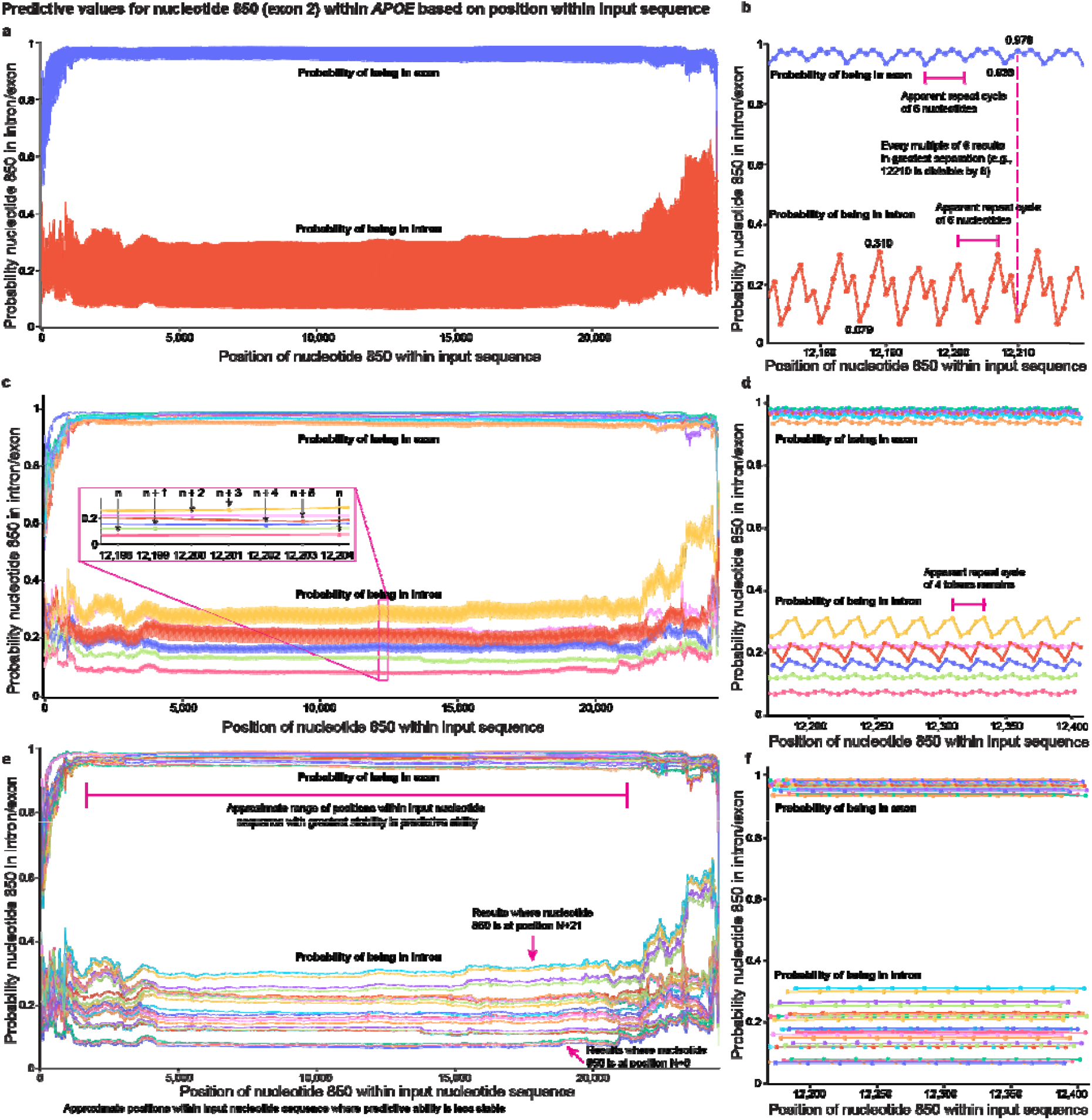
SegmentNT probabilities oscillate in 24-nucleotide intervals. (**a**) SegmentNT probabilities for *APOE* nucleotide 850 (exon 2) for every position within a 24,576-nucleotide input sequence (4096 tokens). Probabilities when nucleotide 850 is towards the middle were more stable for both intronic and exonic predictions but exhibited a clear and surprising oscillating pattern. Both intronic (higher) and exonic (lower) probabilities were worse and less stable towards the outsides of the input sequence. (**b**) A zoomed in plot of figure **a** showing what initially appeared to be a six-nucleotide repeat cycle for both intronic and exonic probabilities for nucleotide 850. Intronic probabilities oscillated between approximately 0.079 (7.9%) and 0.310 (31.0%), while exonic probabilities oscillated between approximately 0.938 (93.8%) and 0.978 (97.8%). Every multiple of six resulted in the greatest separation between the intronic and exonic probabilities (lowest intronic and highest exonic). (**c**) We plotted six sets of values for both intronic and exonic probabilities (every sixth value), which should result in a linear, non-cyclical pattern, but a four-token cycle remained. Colors indicate separate lines resulting from every multiple of six (e.g., n, n+1, etc.). (**d**) Zoomed in plot of **c**. (**e**,**f**) Plotting every 24^th^ value res lts in linear probabilities within the middle portion of the input sequence. Distal positions remained sporadic (indicated by grey shading). Colors indicate separate lines resulting from every multiple of 24. Original interactive plotly HTML files available in **Supplementary Data** for deeper exploration.

Since the nucleotide being assessed remained constant, probabilities should also remain constant, unless position inherently biases the model. Since we previously established that first and last positions resulted in lower and more variable probabilities than middle positions, we therefore expected the tails of the input sequence to be lower and less stable, which we observed (**Fig. 6a**).

Surprisingly, however, intronic and exonic probabilities varied dramatically and cyclically within the central window when shifting nucleotide 850 by only a few positions (**Fig. 6a**,**b**). Specifically, intronic probabilities consistently varied between approximately 0.079 (7.9%) to 0.310 (31.0%; range: 0.068 to 0.311) by shifting nucleotide 850 only three positions (**Fig. 6b**). Exonic probabilities varied from approximately 0.938 (93.8%) to 0.978 (97.8%; range: 0.933 to 0.984) by shifting two positions. On initial inspection, probabilities appeared to oscillate at six-nucleotide intervals (**Fig. 6b**), where every multiple of six resulted in the greatest separation between intronic and exonic probabilities (lowest intronic and highest exonic).

To empirically test whether the bias operated on a six-nucleotide cycle, we plotted nucleotide 850’s probabilities in six different sets (for both exon and intron predictions), where each set plotted every sixth probability (**Fig. 6c,d**). For example, set one (nucleotide n) plotted probabilities when nucleotide 850 was in positions 0, 6, 12, etc. and set two (nucleotide n + 1) plotted probabilities for positions 1, 7, 13, etc. (**Fig. 6c**). If the bias oscillated on a six-nucleotide cycle, plotting every sixth position should result in a linear, non-cyclical pattern. A new pattern oscillating every four tokens emerged, however, suggesting an inherent cyclical bias remained (**Fig. 6c,d**). Notably, lines plotting every n^th^, n + 1, and n + 2 value had the lowest variability (**Fig. 6c,d**). Lines plotting every n^th^ value (divisible by six) produced the best (lowest) intronic probability, while the line plotting every n + 2 value produced the best (highest) exonic probability. We then plotted every 24^th^ position, resulting in non-cyclical probabilities (**Fig. 6e**,**f**), suggesting that the cyclical pattern occurs in 24-nucleotide intervals and further demonstrating position significantly biases SegmentNT’s probabilities. Distal positions resulted in more sporadic probabilities. Results across the other four primary genes using a randomly selected exonic nucleotide were similar (**Supplemental Fig. 22**). We tested five additional nucleotides for each of the five validation genes (**Supplemental Figs. 23-27**), totaling 30 nucleotides across the ten genes, all of which demonstrated strong cyclical behavior across the input sequence. An analysis of variance (ANOVA) showed that the difference in means between the cyclical groups reported *P*≪0.001. Tukey’s honestly significant difference (HSD) post-hoc test showed that all groups were significantly different from most other groups with a few overlapping groups that varied depending on the gene (**Supplemental Table 3**).

### The distance for probabilities to stabilize varies by gene

We next wanted to assess the distance from each end (i.e., the number of positions) required for the probabilities to stabilize. The distance from each end for *APOE*’s nucleotide 850 to reach a rolling average (window size = 6 positions) ≥0.90 and ≥0.95 appeared greater (i.e., “slower”) from the beginning of the input sequence than from the end (**Fig. 6a,c,e**; **Supplemental Table 4**). We found, however, that the distance to stabilize varied by gene and there was no obvious pattern regarding which end of the input sequence stabilized more quickly. For example, from the beginning of the input sequence, the rolling average for *APOE*’s nucleotide 850 did not reach ≥0.90 and ≥0.95 until positions 678 and 1,152, respectively, while it reached ≥0.90 and ≥ 0.95 exactly 78 and 2,203 nucleotides from the end (positions 24,498 and 22,373), respectively. Probabilities for *EGFR* stabilized more quickly and consistently from both ends (distance from beginning to reach ≥ 0.90 = 22; end = 23; distance from beginning to reach ≥ 0.95 = 67; end = 68). Distances for the other genes varied widely (**Supplemental Table 4**). These results further demonstrate that position significantly biases SegmentNT’s probabilities and that it should be accounted for when interpreting results.

## Discussion

SegmentNT and other artificial intelligence models have proven extremely powerful and useful across a range of applications. As the use of these tools continues to grow, they will increasingly be applied to research by non-specialists and researchers in domains outside of computer science. As such, end users and model architects must be aware of potential biases and limitations of these powerful tools.

Here, we demonstrated two intrinsic biases (input sequence length and position, therein) affecting SegmentNT’s raw intron and exon probabilities that may generalize to the other twelve SegmentNT attributes not assessed herein, as well as other nucleotide transformer models. We clarify, however, that position within the input window is not likely an inherent bias in the model itself, but rather due to the lack of sufficient surrounding sequence context. Though, regardless of whether a bias is inherent to the model itself or due to practical use limitations, the bias exists and should be properly managed when interpreting raw output. Additionally, although sequence input length technically biases SegmentNT’s results, this does not represent an inherent problem with the model but demonstrates the biological complexity.

We assessed only two of the 14 features that SegmentNT predicts because intron and exon probabilities were sufficient to demonstrate the biases and to provide a focused analysis of their nature. The biases likely affect the other twelve features as well, but future tests will be necessary to confirm. Thus, we propose that the following should be carefully considered when developing or using nucleotide transformer models: (1) position within the input sequence, and (2) input sequence length.

The nature of the probability distributions was clearly different between the first, middle, and last positions of the input sequence, where first and last exhibited greater variability. It was not surprising that raw values in the middle position appeared to be the most reliable, at face value, since the first and last positions do not benefit from the same amount of sequence context. We were surprised, however, to find that the first and last positions still provided strong predictive value—when properly interpreted and managed. Specifically, when interpreting the raw values, it is not possible to set a single threshold that accurately classifies a nucleotide as intronic or exonic across all three distributions. It is possible, however, to normalize across all three distributions to set a single threshold value. This shows that, while there is an inherent bias in the raw values (based on practical use limitations), the model itself is remarkably robust—even with limited sequence context.

When assessing every position within the input sequence for a given nucleotide (e.g., *APOE* nucleotide 850), and not just the first, middle, and last positions, there was a surprisingly large variance in the probabilities when moving the nucleotide only a few positions in either direction, demonstrating that even small changes in position can have significant and meaningful impacts on a given nucleotide’s prediction. This pattern likely explains the variability observed across a given gene, as shown in Figures 1, 2, and 3. Why there is such large variance within a small span is unclear and merits deeper investigation.

Surprisingly, the variance exhibited a cyclical pattern that initially appeared to oscillate in six-nucleotide intervals but actually occurred in 24-nucleotide intervals. The nature of this cycle and pattern is also unclear and merits deeper investigation. Given that the cycle occurs in a multiple of six, it could be related to SegmentNT’s base model having been trained on six-nucleotide tokens with the U-Net architecture requiring 2^n^ tokens where n≥2 (i.e., minimum number of nucleotides is 6 nucleotides per token × 2^2^ tokens = 24 nucleotides). Alternatively, the oscillation could occur from the NTK-aware relative positional embeddings (RoPE) that have a periodic nature^38,51^ and similarly require 2 × n tokens (at least 4 tokens, or 24 nucleotides). Retraining the Nucleotide Transformer with different-sized tokens may provide additional insights^52,53^. In principle, one method to address this cyclical variation would be to average the probabilities across a sliding window, but this would dramatically increase the computational costs because it would require running SegmentNT multiple times. Such an approach would also only be a superficial fix to an underlying variation that should be addressed in the model itself.

As expected, the length of the input sequence significantly impacts the model’s performance simply because having greater sequence context provides more information to model, but the input size required to obtain most of the benefit was shorter than expected at only 3,072 nucleotides in this study across ten genes. This finding suggests that increasing the context size beyond ∼3,072 nucleotides may significantly increase computational load without producing meaningful gains in model performance. We also presented data suggesting SegmentNT’s probabilities may capture additional biological information (e.g., alternative splicing), though these results were purely observational and further work is needed.

This work includes several important limitations that must be considered. One limitation is that we included analyzed five genes and a single non-genic region (negative control) plus five validation genes. Though the results were largely consistent, they may not generalize to the entire genome. We were limited in the number of regions we could include because of the computational resources required to place every nucleotide at the first, middle, and last position. Despite this limitation, our results are likely a strong representation of most genes since the behavior was consistent across all regions.

Another limitation is that we also cannot state at this point whether discrepancies between SegmentNT’s probabilities and the gene annotation are due to inaccuracies in SegmentNT or because of inaccurate gene annotations. We used canonical RefSeq exons as the truth set when assessing SegmentNT’s performance, but these annotations remain imperfect^54,55^, as demonstrated by several examples where both SegmentNT and reported Ensembl RNA isoforms contradicted RefSeq. It is notable, however, that some of the seemingly discrepant *EGFR* exonic predictions from SegmentNT were, in fact, correct. Multiple peaks (indicated by # in **Fig. 3a**) corresponded to *EGFR-AS1* exons on the opposite strand, highlighting that end users must consider both DNA strands when using SegmentNT and not only interpret results from a single gene’s or strand’s perspective. Indeed, this is one of the primary hopes and uses of artificial intelligence and machine learning models such as SegmentNT. As suggested by SegmentNT’s authors, it is meant to reveal new insights into a genome, including annotating *de novo* genomes^38^. Additional work will be needed to systematically vet discrepancies between SegmentNT and existing annotations.

A third limitation is that, even though we demonstrated the variation in SegmentNT’s raw probabilities across every position in a 24,576-nucleotide input sequence using 30 individual nucleotides across ten genes (e.g., *APOE*’s nucleotide 850), this merely assesses probabilities for a small number of nucleotides. Much larger studies may be necessary to establish a complete picture. We also presented two simplistic methods for normalizing probabilities across the input sequence, but more sophisticated solutions may be able to elucidate additional information.

Despite these limitations, we demonstrated two major biases affecting SegmentNT that may generalize across other foundation models implementing similar architectures. Future use cases attempting to leverage such models should account for these biases to ensure that results are consistent and interpreted properly. These issues can compound with each layer of complexity, potentially causing large downstream effects. Accounting for these, and other unmeasured biases, will be essential in advanced use cases such as predicting variant-level effects on pathogenicity or leveraging nucleotide transformers in a clinical setting.

Our study clearly describes innate biases in SegmentNT’s raw probabilities that may generalize across other nucleotide transformer models and provides actionable and generalizable insights. We hope our study contributes to making SegmentNT and other models even better and that this information will help mitigate these issues in future work, improving the reliability of nucleotide-resolution annotations in genomic LLMs.

## Methods

### GitHub repository and example code

Step-by-step instructions and example code for replicating these results are publicly available at https://github.com/jmillerlab/nt_context_evaluation. Additional descriptions of the methods are outlined below.

### Compute environment

All analyses were conducted on a high-performance computing cluster running Ubuntu 20.04 LTS. The cluster is equipped with four NVIDIA A100-SXM4-40GB GPUs that are interconnected via NVLink, enabling high-speed communication between them. The system utilizes CUDA 12.6, and the software environment includes Python 3.10.12 along with essential libraries such as argparse, haiku, jax, json, matplotlib, nucleotide_transformer.pretrained^39^, numpy, os, pandas, plotly^56^, scipy, seaborn, sklearn, statistics, and sys. The nucleotide_transformer_pretrained module^39^ is available at https://github.com/instadeepai/nucleotide-transformer and contains the pretrained SegmentNT^38^ model parameters that were used to calculate nucleotide-level exon and intron predictions.

Compute nodes featured 96 CPUs powered by AMD EPYC 7713 24-Core processors with a base clock speed of 2.65 GHz. The system is configured with 8 non-uniform memory access (NUMA) nodes and 500GB of DDR4 RAM, providing sufficient resources for preprocessing and data loading tasks to ensure smooth handling of these large datasets.

### Reference genome, gene, and variant selection

All tests used the same reference genome that was originally used to train SegmentNT: GCF_000001405.26 (GRCh38 version 26; released August 27, 2024) and downloaded from https://ftp.ncbi.nlm.nih.gov/genomes/all/GCF/000/001/405/GCF_000001405.26_GRCh38/ in January 2025. When selecting genes to assess in this study, for the first set of genes presented in main text results, we selected the five most-studied genes in the human genome as of 2017^40^, which include (in order): *TP53, TNF, EGFR, VEGFA*, and *APOE*. The genes selected for additional validation (*DPM2, ECM1, LINC00207, NAV2-AS5*, and *WFDC5*) had to meet the following criteria: (1) be located on a primary chromosomal contig (e.g., chr1, chr2, etc.); (2) be annotated as a “gene” or a “lncRNA”; (3) have a length ≥ 3,000 and < 10,000 (to avoid longer genes because of computational resources); (4) not be pseudogenes or genes lacking standard gene symbols (i.e., omitted genes beginning with “LOC”); and (5) be located on a different chromosome than any of the original five genes selected (to test across additional chromosomes). We also required each additional gene to be on different chromosomes from each other, and two had to be from chromosomes not included as part of the Nucleotide Transformer’s^39^ training set (i.e., they had to be from chromosomes 20-22). For the ten genic regions assessed, all ten are uniquely mappable using short-reads, demonstrating that they consist of unique (i.e., unambiguous) sequence because they are not camouflaged, per our^57,58^ and others’^59^ previous work; having significant repeat sequence could bias the results. Only *TNF* appears to be in a genomic region that is difficult to resolve; it falls within the major histocompatibility complex on chromosome six, which is known to have many alternate haplotypes, but the gene itself is not repetitive.

The non-genic control region was selected to have similar characteristics to the *APOE* gene in size and repeat content. Using Python’s random module, we selected 20 random sequences from the reference genome by randomly selecting a chromosome and starting position, making the regions the same size as *APOE* (3,612 nucleotides based on RefSeq). The selected regions were restricted to non-genic, non-centromeric, and non-telomeric regions. We then used RepeatMasker (version 4.1.5) to quantify the amount of repeat content—short interspersed nuclear elements (SINEs), long interspersed nuclear elements (LINEs), and long terminal repeats (LTRs)—in *APOE* (chr19:44905782-44909393; 19% SINEs, 0% LINEs, 0% LTRs) and in the randomly selected genomic regions (**Supplemental Table 5**). We selected chr3:153050001-153053613 because it has similar repeat content (24% SINEs, 0% LINEs, 0% LTRs).

To assess the stability of probabilities across the full window of a 24,576-nucleotide input sequence, we selected nucleotides from each gene to plot their probabilities across the full range. For the five primary genes in our analyses, we randomly selected an exonic variant. For the five validation genes, we selected five variants across the gene, including deep within exons and at exon boundaries.

RefSeq gene annotations (exon coordinates, specifically) were extracted from the accompanying General Feature Format (gff) file for each genomic region using the gene field supplied in the gff file. We extracted the gene regions and included 50k bases upstream and downstream of each gene using the `samtools faidx` command. The 50k padding was used to evaluate the effects of nucleotide context on nucleotide-level predictions since the sequence before and after the gene was used to make predictions. The length of the padding sequence can be changed depending on the context window length. For RNA isoform structures, we used version 113 of the Ensembl annotations for the same reference genome.

### Running SegmentNT

SegmentNT (30kb training context using the Nucleotide Transformer backbone) was executed using a sliding window of 1 nucleotide to iterate over all possible contexts for each gene sequence using our custom script `run_nt.py`. When assessing nucleotide position bias, we used a static window size of 24,576 nucleotides (**Fig. 1**). To assess input size bias, we used eleven window sizes of 2^n^ tokens where 2 ≤ n ≤ 12 and each token is 6 nucleotides. Thus, window sizes ranged from 6 × 2^2^ tokens = 24 nucleotides to 6 × 2^12^ tokens = 24,576 nucleotides). The default SegmentNT output was then parsed and sorted to facilitate locating predictions at any position within the input sequence using `parseWindow.py`. The parsed output file contains lists of all intron and exon predictions sorted by position within the context window, starting with a list of probabilities for each position within a gene where the nucleotide of interest was the first nucleotide in the input sequence and ending with a list of predicted probabilities where the nucleotide of interest was the last nucleotide in a prediction window. That parsed file is then passed to two separate scripts: (1) `getPositions.py`, which extracts SegmentNT probabilities for the first, middle, and last predictions; and (2) `getOnePosition.py`, which extracts all probabilities at all contexts for a specific nucleotide within a gene.

### Statistical analyses and graphing

We provide several additional scripts on our GitHub repository to analyze and graph these results. The `graph_diff_Exon_Intron.py` script generates five output files: (1) A plot graphing the area under the curve (AUC) calculated for the first, middle, and last positions; (2) A statistics metadata file for the last position in the context window including the average ± standard deviation (median) exon predictions in exons and intron, true positive, false positive, false negative, true negative, sensitivity, specificity, and accuracy; (3) A statistics metadata file for the middle position in the context window containing the same statistics as above; (4) A statistics metadata file for the first position in the context window containing the same set of statistics as the last and middle predictions; and (5) An HTML file showing the difference in exon and intron prediction across the entire gene. Those results were then used to generate the graphs and tables included in this manuscript. We used NCBI RefSeq exons as the gold standard (“truth”) when assessing SegmentNT’s model performance (accuracy, sensitivity, and specificity).

We also include a script, `graph_change_in_predictions_based_on_length.py`, to graph how context length impacts predictions. Similarly, this script plots the AUC values from different context lengths and allows users to normalize the results through Z-score normalization, subtracting the intron probability from the exon probability, or using the raw unnormalized values.

The `graph_pred_at_position.py` script is used to visualize the impact of nucleotide position within the input sequence on predicted exon or intron probabilities. That script looks at a single nucleotide within a gene and generates an HTML graphing how the prediction changes based on nucleotide position within the sequence. Additionally, the `graph_different_contexts.py` script was used to graph exon/intron probabilities across each gene using the first, middle, or last nucleotide within a context window.

When comparing SegmentNT’s probability distributions between first, middle, and last positions, we used a t-test; based on the large sample size and the Central Limit Theorem, this is reliable even if the distributions do not follow a perfect normal distribution. Wherever Python reported a p-value of 0.0, we reported the p-value as being < 2.23e-308. Reporting a p-value of 0.0 is inaccurate and 2.23e-308 is the current lower limit of Python’s positive float value, based on Python’s (v3.10.12) value of `sys.float_info.min`.

We calculated statistical differences between predictions based on different nucleotide contexts within a context window to determine the extent to which oscillation impacts predictions using the `calculateANOVAandTukey.py` script. An ANOVA was calculated using all 24 groups of expected oscillating predictions. Then Tukey HSD was used to calculate pairwise differences between each of the 24 groups. We opted to remove ∼20% from either tail (4,800 contexts) because of visually high variability when there is relatively little context either before or after the nucleotide of interest.

### Isoforms and expression plots

Isoform expression plots based on human frontal cortex data were generated by data from Aguzzoli-Heberle et al.^45^ at https://ebbertlab.com/brain_rna_isoform_seq.html. Isoform expression plots based on nine GTEx samples were generated at https://ebbertlab.com/gtex_rna_isoform_seq.html; these data were generated by Glinos et al.^47^ and analyzed by Page et al.^46^. Isoform structures throughout the manuscript were plotted using RNAPysoforms^48^. In the original paper by Aguzzoli Heberle et al.^45^, the minimum expression criteria to be considered “expressed” was a median counts-per-million (CPM) ≥ 1 across all samples. Thus, here, if an isoform did not reach this threshold, it could not be considered as being the most expressed isoform (see Figure 3).

## Data Availability

All scripts used to conduct these analyses are publicly available and can be found at https://github.com/jmillerlab/nt_context_evaluation. A selection of results is included on the GitHub repository and through the Supplementary Data located at https://zenodo.org/records/15025141. However, some of the intermediate output files were exceptionally large (>500G). The commands used to generate those files are publicly available, and the files themselves are also available upon reasonable request.

## Acknowledgments

This work was supported by the BrightFocus Foundation and its donors [A2020118F to Miller; A2020161S to Ebbert], the National Institutes of Health [1P30AG072946-01 to the University of Kentucky Alzheimer’s Disease Research Center; AG068331 to Ebbert; R35GM138636 to Ebbert; R01AG08273 to Fardo; RF1AG082339 to Fardo], and the Alzheimer’s Association [2019-AARG-644082 to Ebbert]; PhRMA Foundation Predoctoral Drug Discovery Fellowship to Heberle; and NSF MRI grant 2216140 (Co-PI Bumgardner). The content is solely the responsibility of the authors and does not necessarily represent the official views of the National Institutes of Health.

