## Supplementary Figures for "Systematic contextual biases in SegmentNT potentially relevant to other nucleotide transformer models"

Mark T. W. Ebbert^1,2,3,^,*^, Anna Ho^2,4,^^, Madeline L. Page^1,2,3,^^, Bram Dutch^5^, Blake K. Byer^2,4^, Kristen L. Hankins^6^, Hady Sabra^1,6,8^, Bernardo Aguzzoli Heberle^1,3^, Mark E. Wadsworth^1,2,3^, Grant A. Fox^1,3^, Bikram Karki^2,5^, Caylin Hickey^4,6^, David W. Fardo^1,7^, Cody Bumgardner^2,4,6^, Yasminka A. Jakubek^1,2,9^, Cody J. Steely^1,2,9^, Justin B. Miller^1,2,4,8,*^

^1^Sanders-Brown Center on Aging, University of Kentucky, Lexington, KY 40506, USA

^2^Division of Biomedical Informatics, Department of Internal Medicine, University of Kentucky, Lexington, KY 40506, USA

^3^Department of Neuroscience, University of Kentucky, Lexington, KY 40506, USA

^4^Department of Pathology and Laboratory Medicine, University of Kentucky, Lexington, KY 40506, USA

^5^Department of Computer Science, University of Kentucky, Lexington, KY 40506, USA

^6^Institute for Biomedical Informatics and the Center for Applied Artificial Intelligence, College of Medicine, University of Kentucky, Lexington, KY 40506, USA

^7^Department of Biostatistics, University of Kentucky, Lexington, KY 40506, USA

^8^Department of Microbiology, Immunology, and Molecular Genetics, University of Kentucky, Lexington, KY 40506, USA

^9^Markey Cancer Center, University of Kentucky, Lexington, KY 40506, USA

Author Disclosures: None

^^^These authors contributed equally and share co-first authorship.

^*^To whom correspondence should be addressed:

Mark Ebbert

Justin Miller

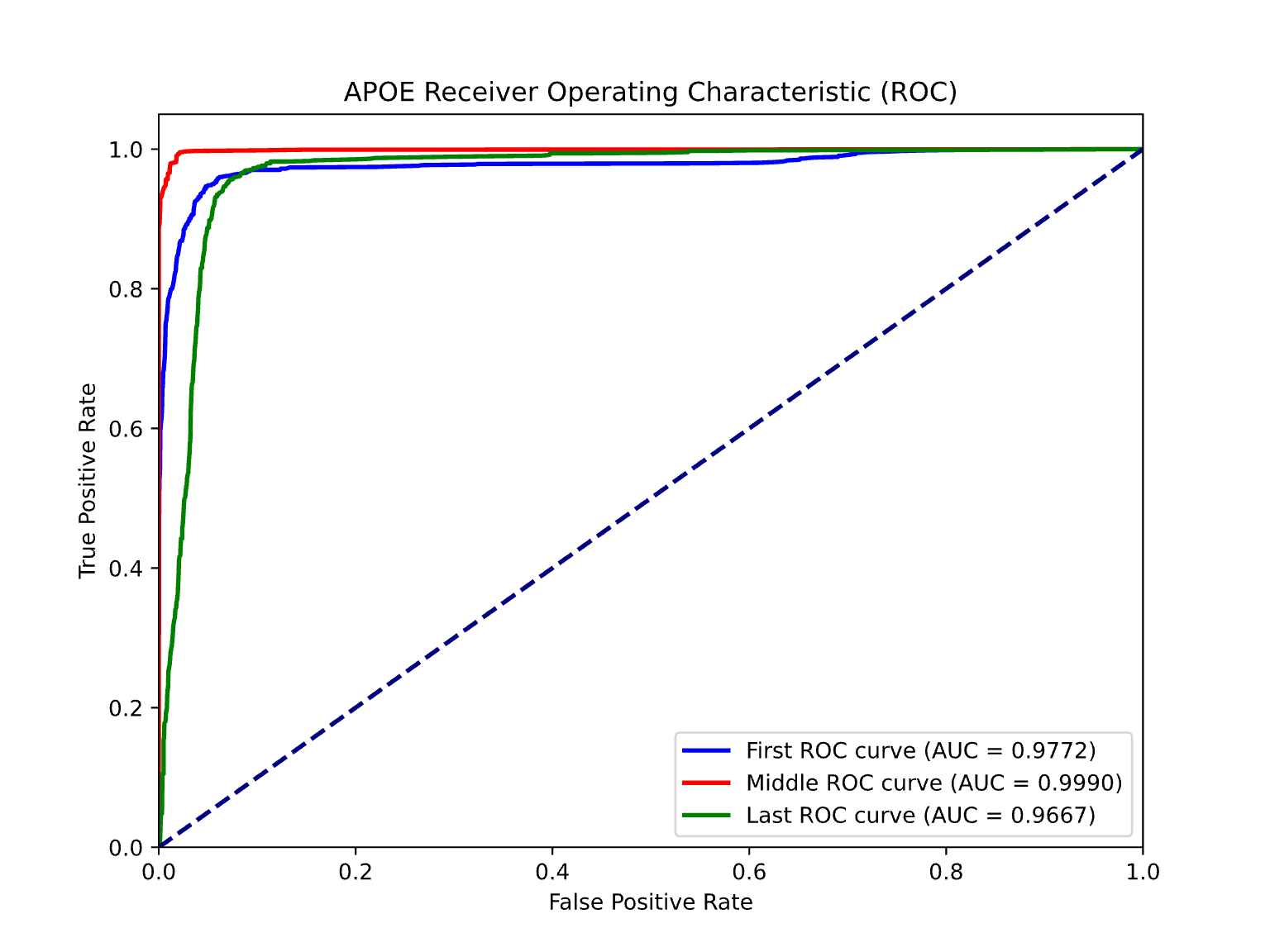

Supplemental Fig. 1. Receiver Operating Characteristic (ROC) curve for raw SegmentNT *APOE* exonic probabilities. Raw probabilities being in the middle of the input sequence perform best, compared to being either first or last in the input sequence. Despite first and last positions appearing to offer low predictive value, when taking raw scores at face value, all three positions provide high predictive value; their individual interpretations would be different given the large difference in scales, however (see main Fig. 2a).

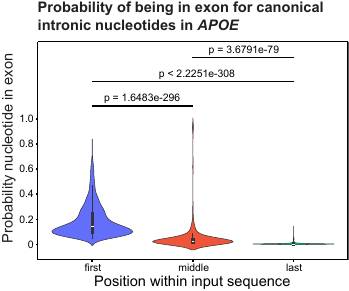

Supplemental Fig. 2. The probability of being in an exon for canonical intronic nucleotides in *APOE*. For nucleotides within canonical introns, the probabilities for being in an exon when the nucleotide was in the middle were significantly lower than being in the first (p = 1.65e-296), but higher than last (p = 3.68e-79) positions.

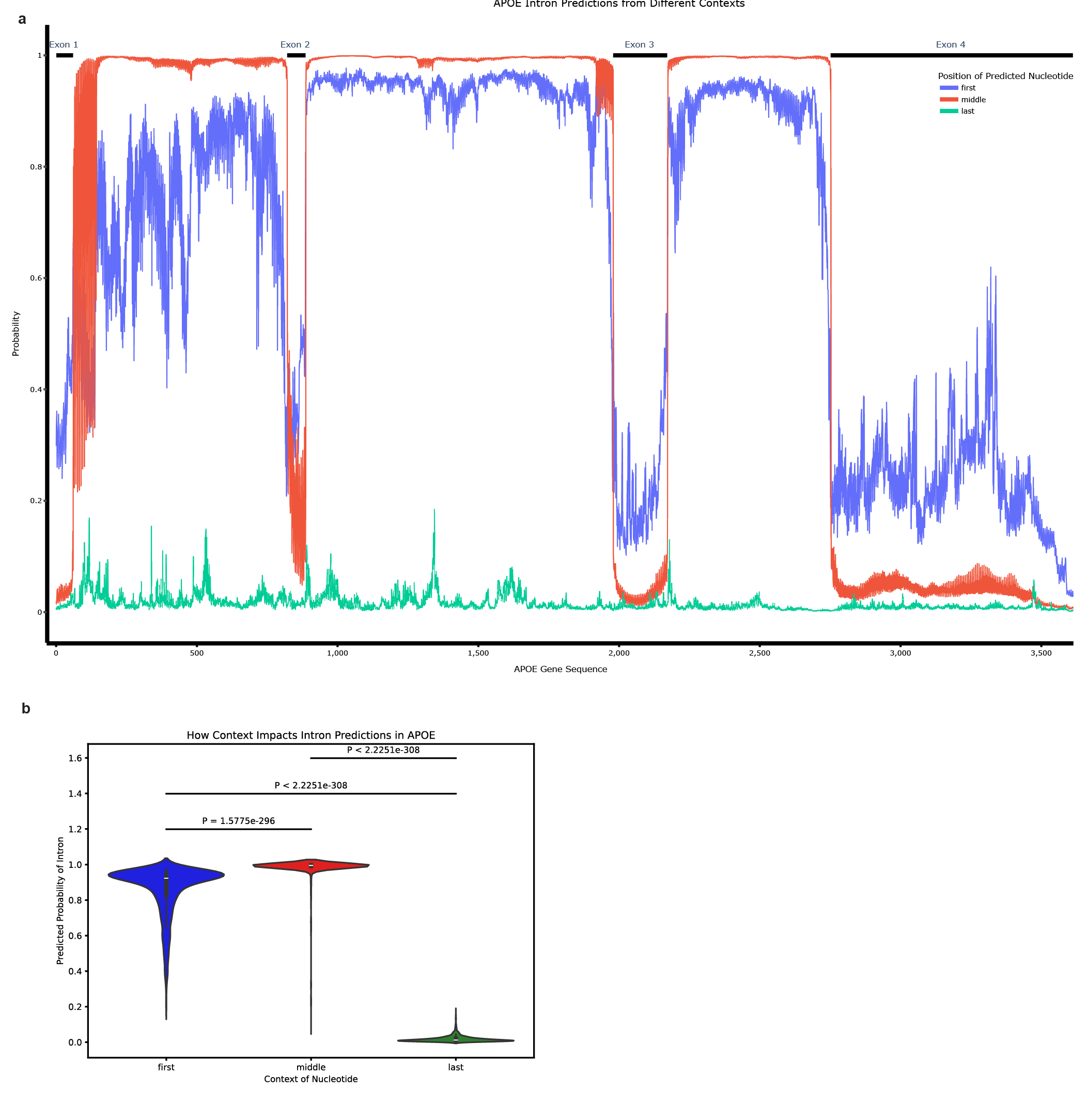

Supplemental Fig. 3. SegmentNT intronic predictions in *APOE*. (**a**) SegmentNT intronic probabilities for each nucleotide in across *APOE*. (**b**) Violin plots showing distribution of probabilities of being in an intron for nucleotides within *APOE* RefSeq introns. These results closely mirror results from SegmentNT’s exonic predictions.

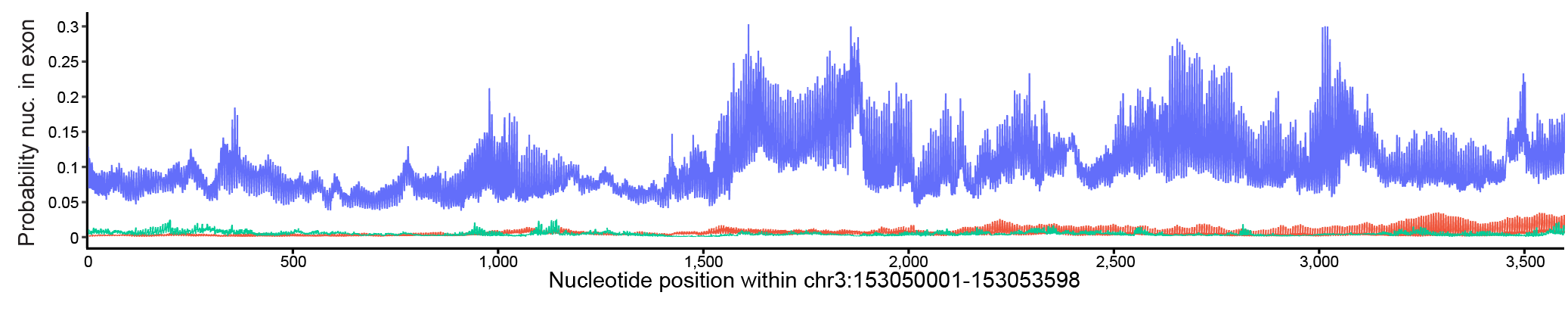

Supplemental Fig. 4. SegmentNT’s exonic probabilities across non-genic negative control region (chr3:153050001-153053598). As a negative control, we tested SegmentNT’s exonic predictions across a non-genic region. As expected, probabilities for all positions in the input sequence were low throughout the region, indicating zero exons. Note that the y-axis scale maxes at 0.3.

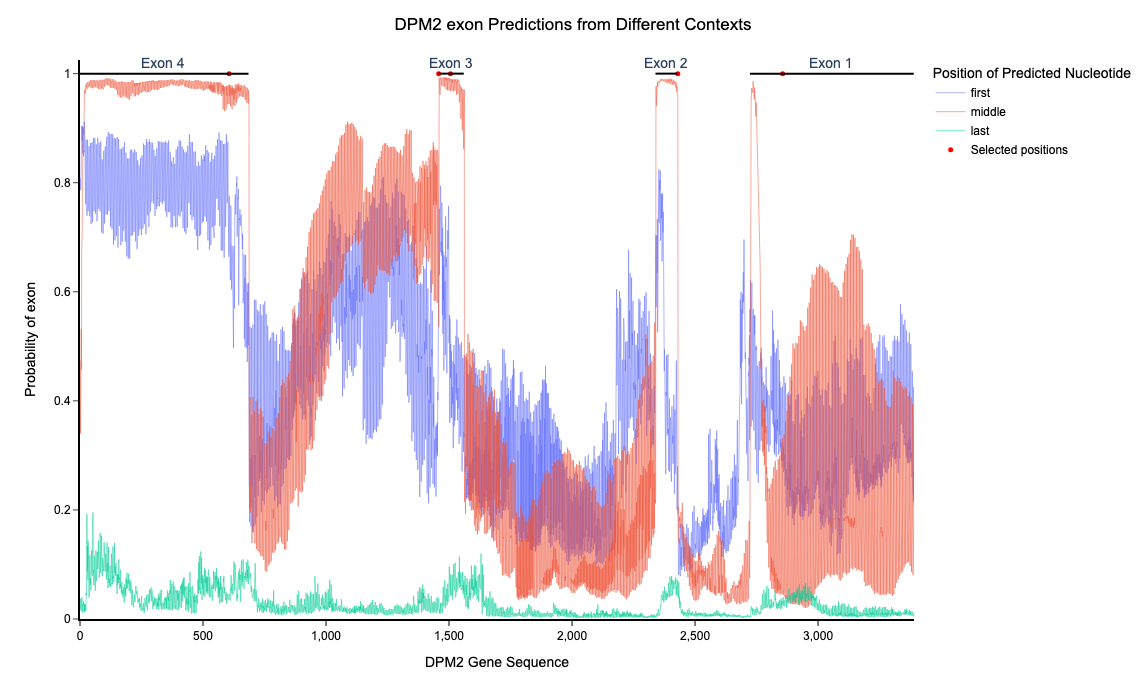

Supplemental Fig. 5. SegmentNT’s exonic probabilities across *DPM2*. SegmentNT probabilities of being in an exon for *DPM2*, along with RefSeq exons (black lines). Red dots indicate exonic nucleotides later tested at every position in the input sequence. Positions include 606, 1458, 1506, 2430, and 2856.

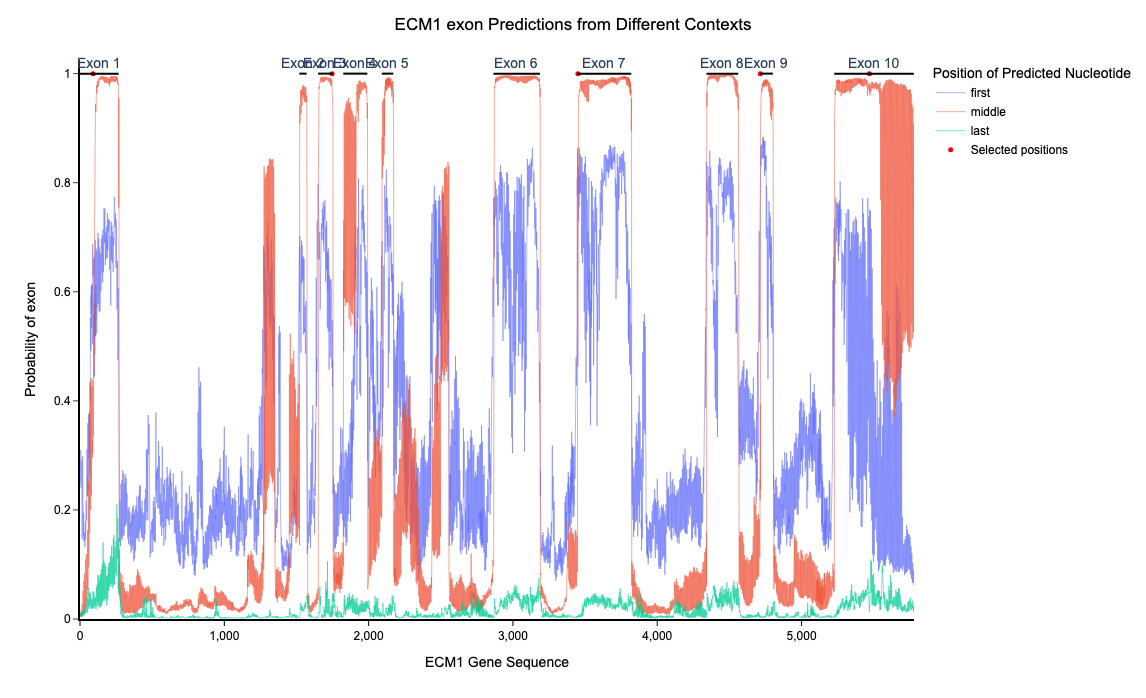

Supplemental Fig. 6. SegmentNT’s exonic probabilities across *ECM1*. SegmentNT probabilities of being in an exon for *ECM1*, along with RefSeq exons (black lines). Red dots indicate exonic nucleotides later tested at every position in the input sequence. Positions include 90, 1747, 3450, 4714, and 5470.

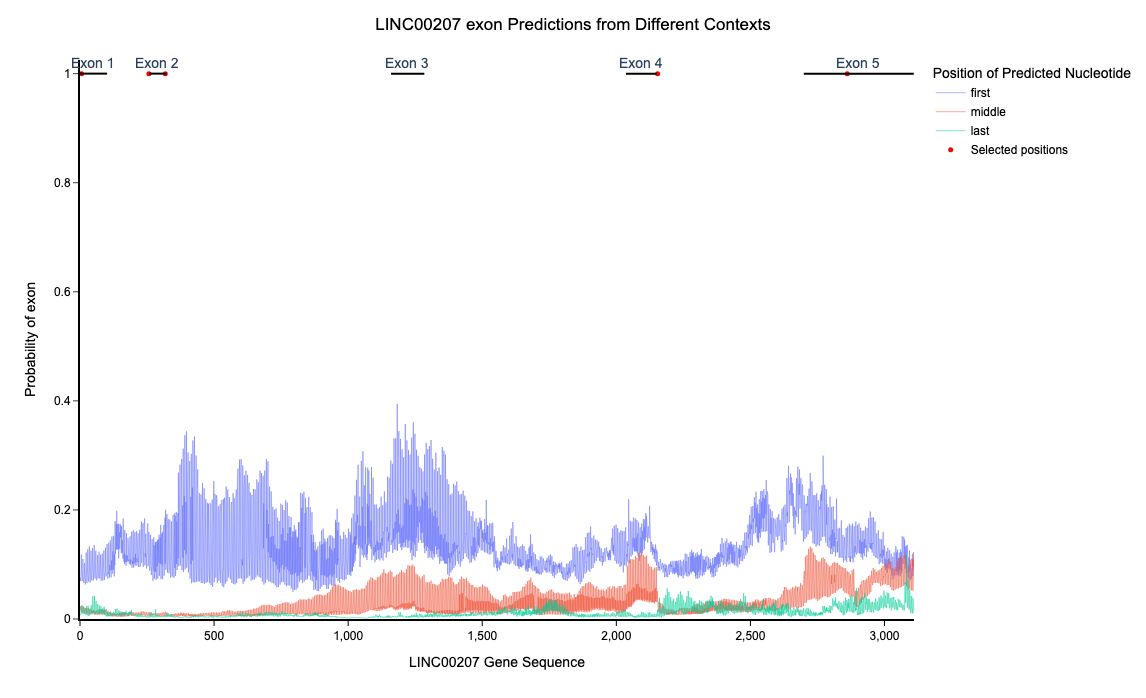

Supplemental Fig. 7. SegmentNT’s exonic probabilities across *LINC00207*. SegmentNT probabilities of being in an exon for *LINC00207*, along with RefSeq exons (black lines). Red dots indicate exonic nucleotides later tested at every position in the input sequence. Positions include 6, 257, 318, 2154, and 2861.

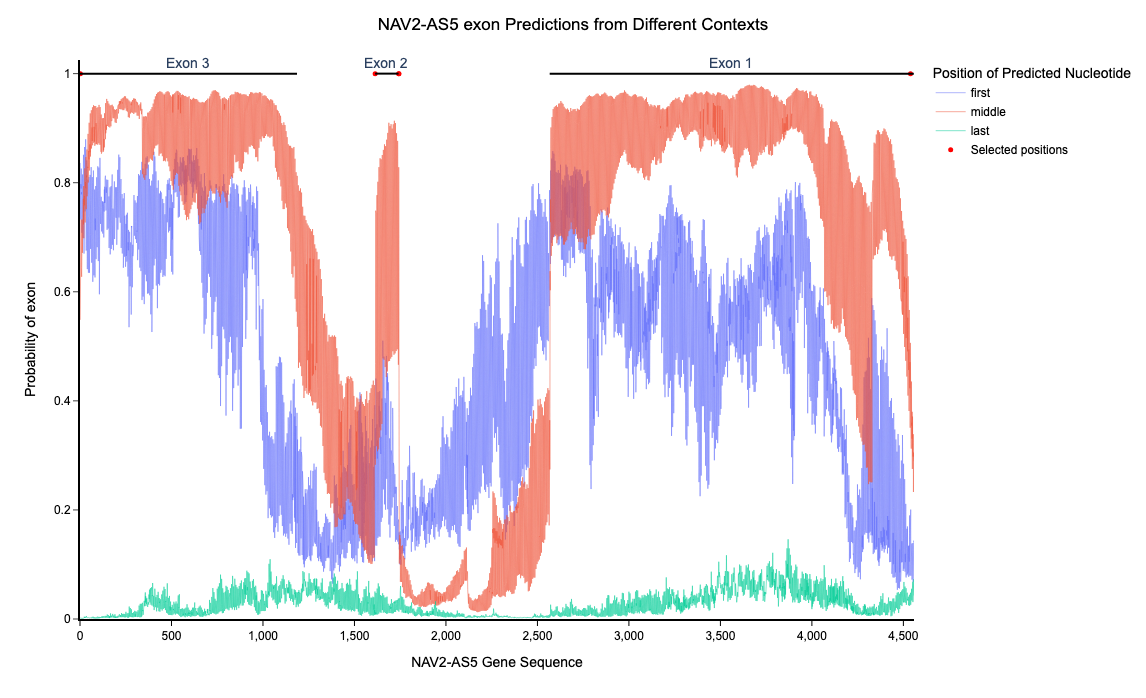

Supplemental Fig. 8. SegmentNT’s exonic probabilities across *NAV2-AS5*. SegmentNT probabilities of being in an exon for *NAV2-AS5*, along with RefSeq exons (black lines). Red dots indicate exonic nucleotides later tested at every position in the input sequence. Positions include 2, 1613, 1742, 1743, and 4539.

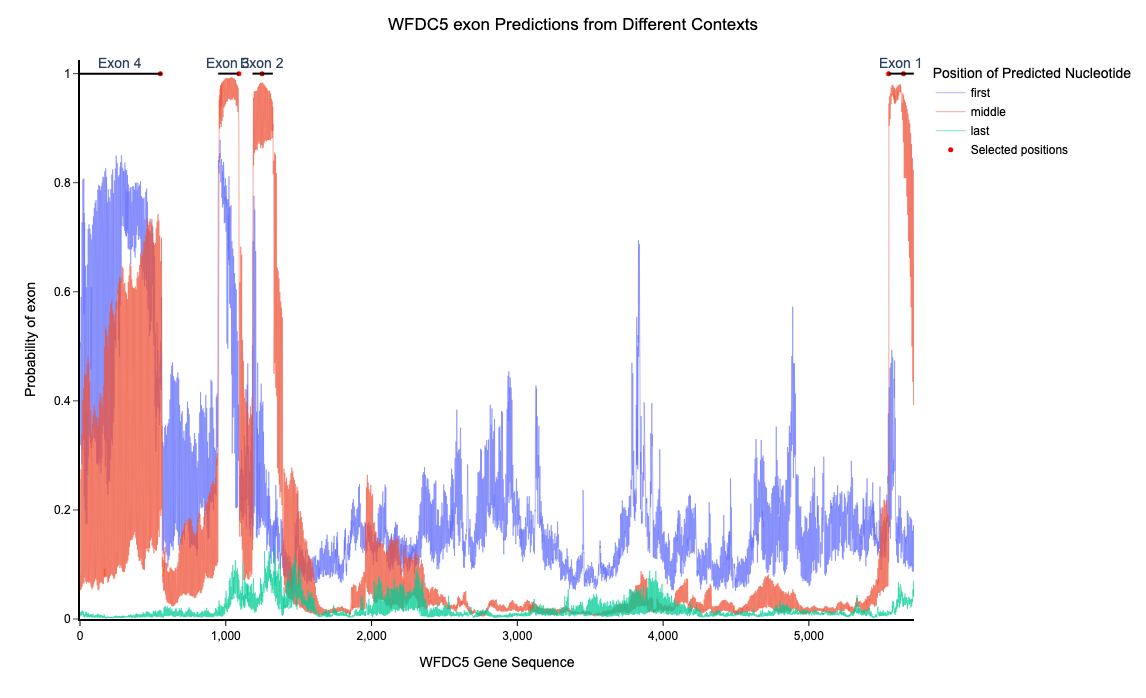

Supplemental Fig. 9. SegmentNT’s exonic probabilities across *WFDC5*. SegmentNT probabilities of being in an exon for *WFDC5*, along with RefSeq exons (black lines). Red dots indicate exonic nucleotides later tested at every position in the input sequence. Positions include 551, 1090, 1249, 5547, and 5649.

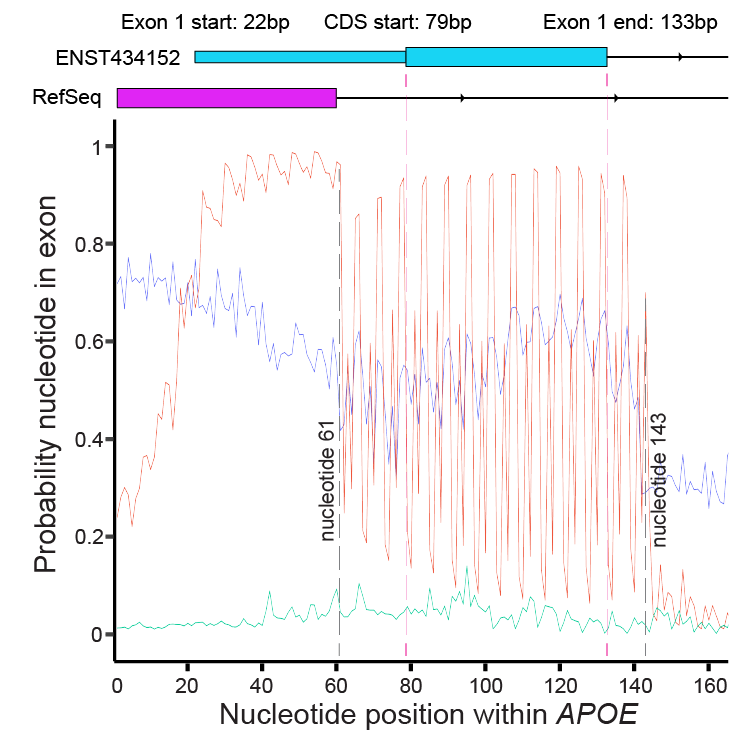

Supplemental Fig. 10. SegmentNT probabilities indicate a strong but inconsistent signal that aligns with extension of exon 1. SegmentNT’s exonic probabilities (using the middle position) indicate a strong but inconsistent signal between approximately nucleotide 61 and 143. This signal closely corresponds with the non-canonical extension of exon 1, per Ensembl *APOE* isoform ENST00000434152, which extends from nucleotide 61 to 133. SegmentNT’s probabilities for this region oscillate dramatically within four nucleotides, ranging from approximately 0.06 to 0.96. Why this signal varies so dramatically is unclear. This plot is a zoomed-in version of main Figure 2a and thus is based on an input sequence of 24,576 nucleotide input sequences (4,096 tokens of six nucleotides).

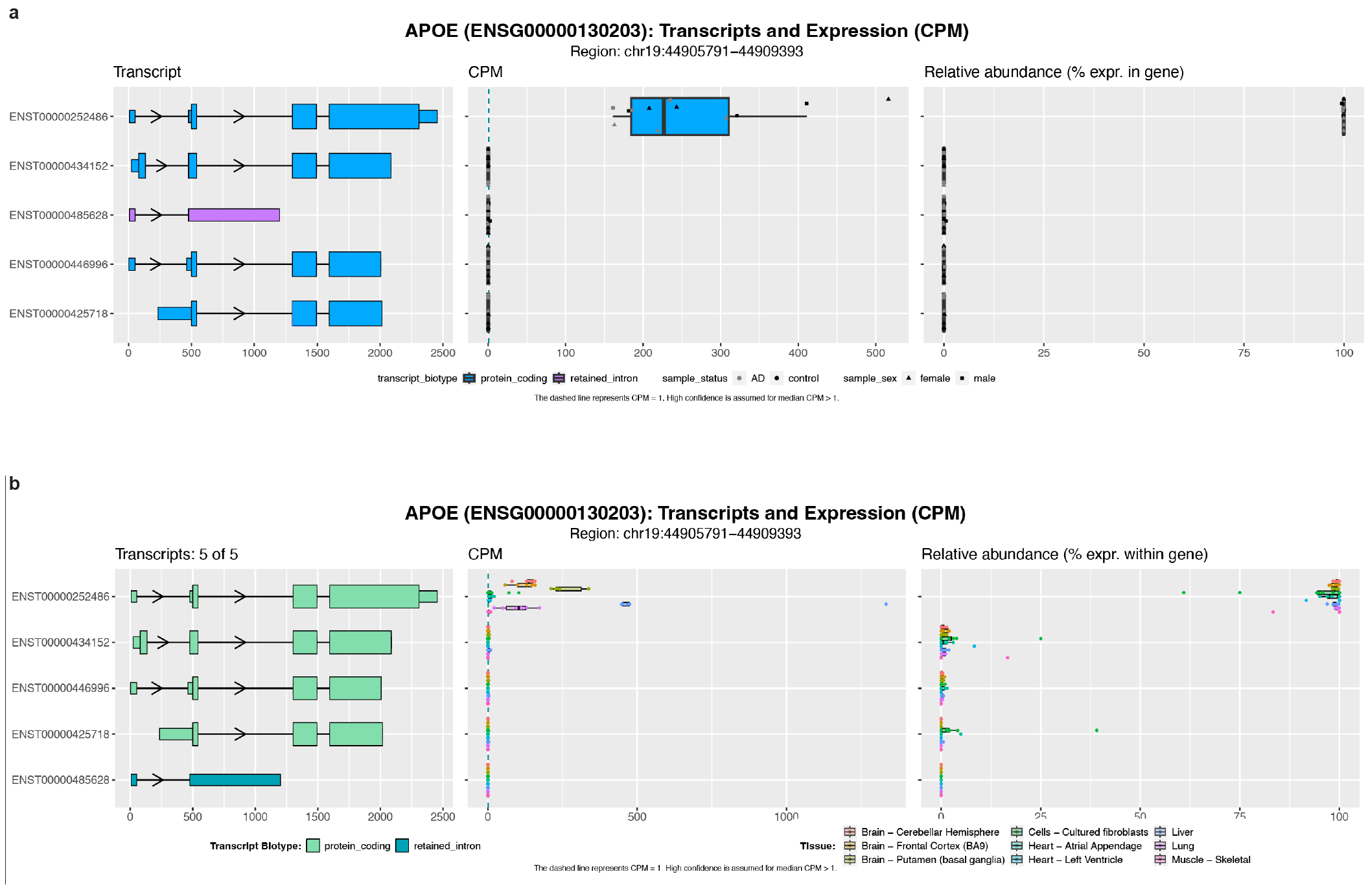

Supplemental Fig. 11. *APOE* isoform expression. (**a**) The top *APOE* isoform expressed in human frontal cortex, per Aguzzoli-Heberle et al. was ENST00000252486. The other four reported isoforms did not have significant expression levels. (**b**) The same *APOE* isoform was the most expressed across nine GTEx tissues, per Glinos et al., based on analyses by Page et al. Isoform expression plots based on human frontal cortex data generated by Aguzzoli-Heberle et al. were generated at <https://ebbertlab.com/brain_rna_isoform_seq.html>. Isoform expression plots based on nine GTEx samples were generated at <https://ebbertlab.com/gtex_rna_isoform_seq.html>; these data were generated by Glinos et al. and analyzed by Page et al.

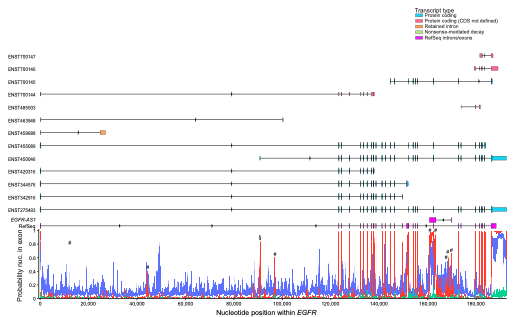

Supplemental Fig. 12. *EGFR* RNA isoforms and SegmentNT exonic probabilities. SegmentNT probabilities of being in an exon for *EGFR*, along with RefSeq exons and all 13 reported Ensembl RNA isoforms. RefSeq exons for *EGFR-AS1* (opposite strand), which align with some of SegmentNT’s exonic predictions, are also included.

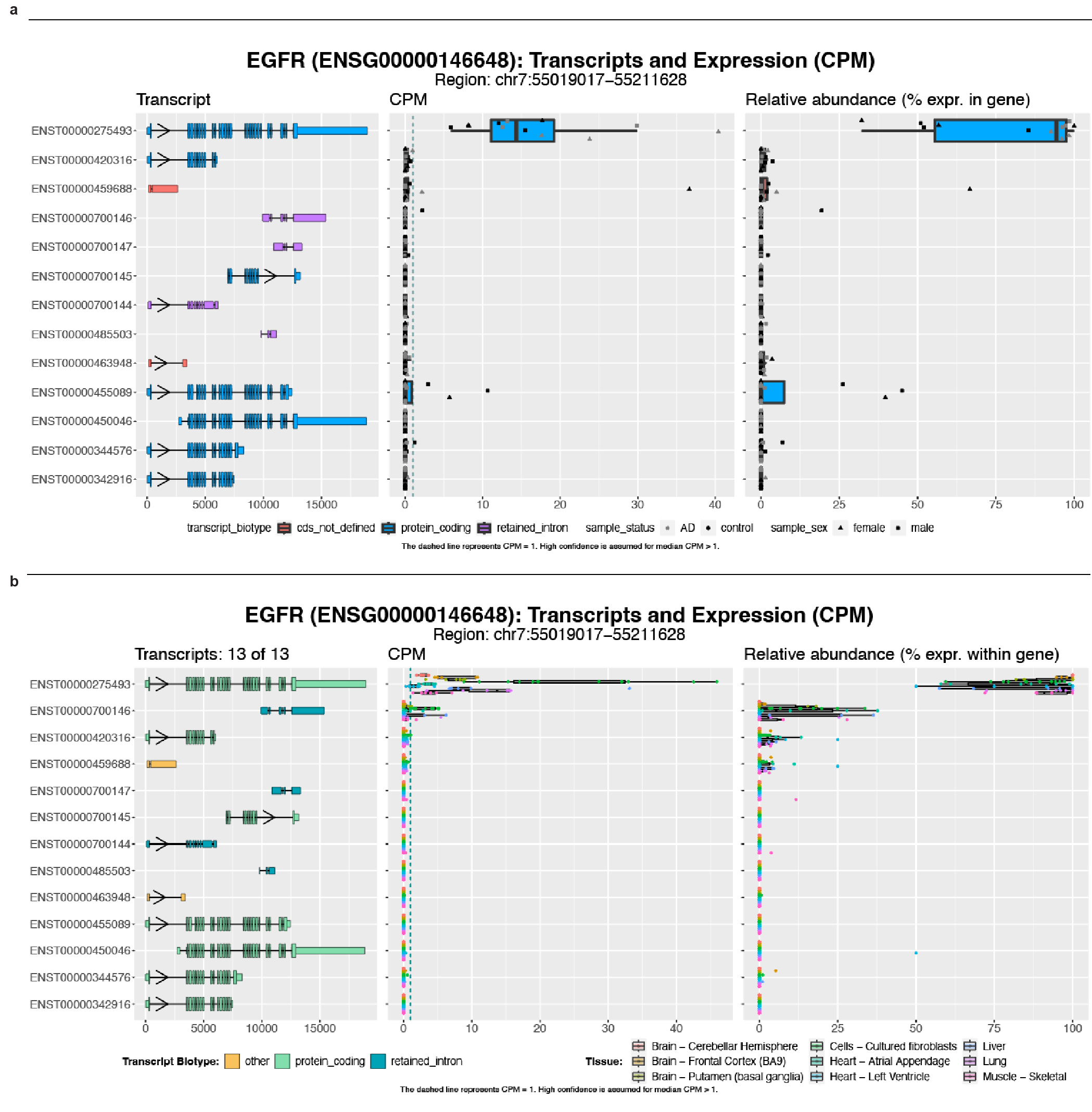

Supplemental Fig. 13. EGFR RNA isoform expression. (**a**) The top *EGFR* isoform expressed in human frontal cortex, per Aguzzoli-Heberle et al. was ENST00000275493. The other reported isoforms did not have significant expression levels. (**b**) The same isoform was the most expressed across nine GTEx tissues, per Glinos et al., based on analyses by Page et al. Isoform expression plots based on human frontal cortex data generated by Aguzzoli-Heberle et al. were generated at <https://ebbertlab.com/brain_rna_isoform_seq.html>. Isoform expression plots based on nine GTEx samples were generated at <https://ebbertlab.com/gtex_rna_isoform_seq.html>; these data were generated by Glinos et al. and analyzed by Page et al.

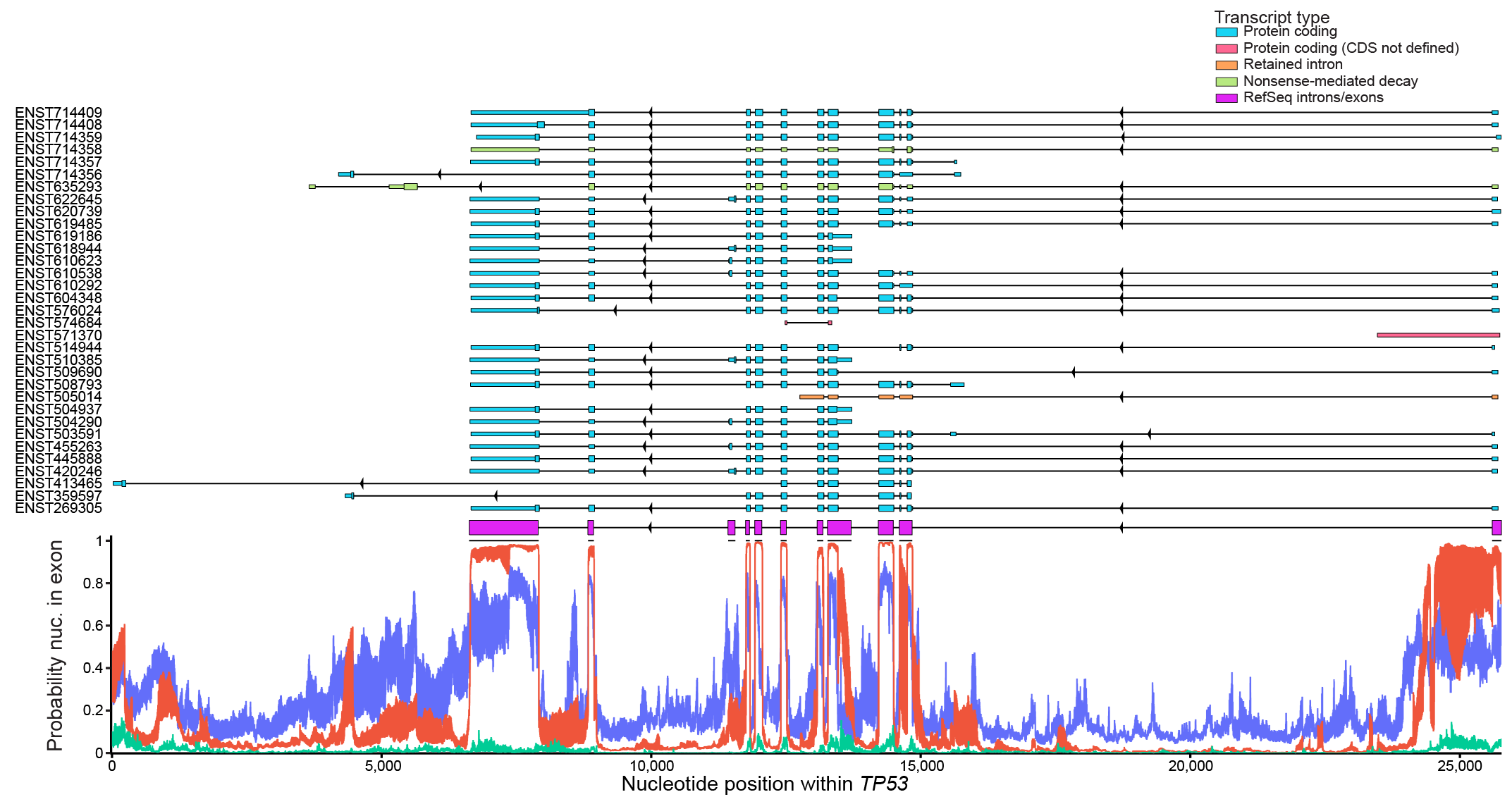

Supplemental Fig. 14. TP53 RNA isoforms and SegmentNT exonic probabilities. SegmentNT probabilities of being in an exon for *TP53*, along with RefSeq exons and all 33 reported Ensembl RNA isoforms.

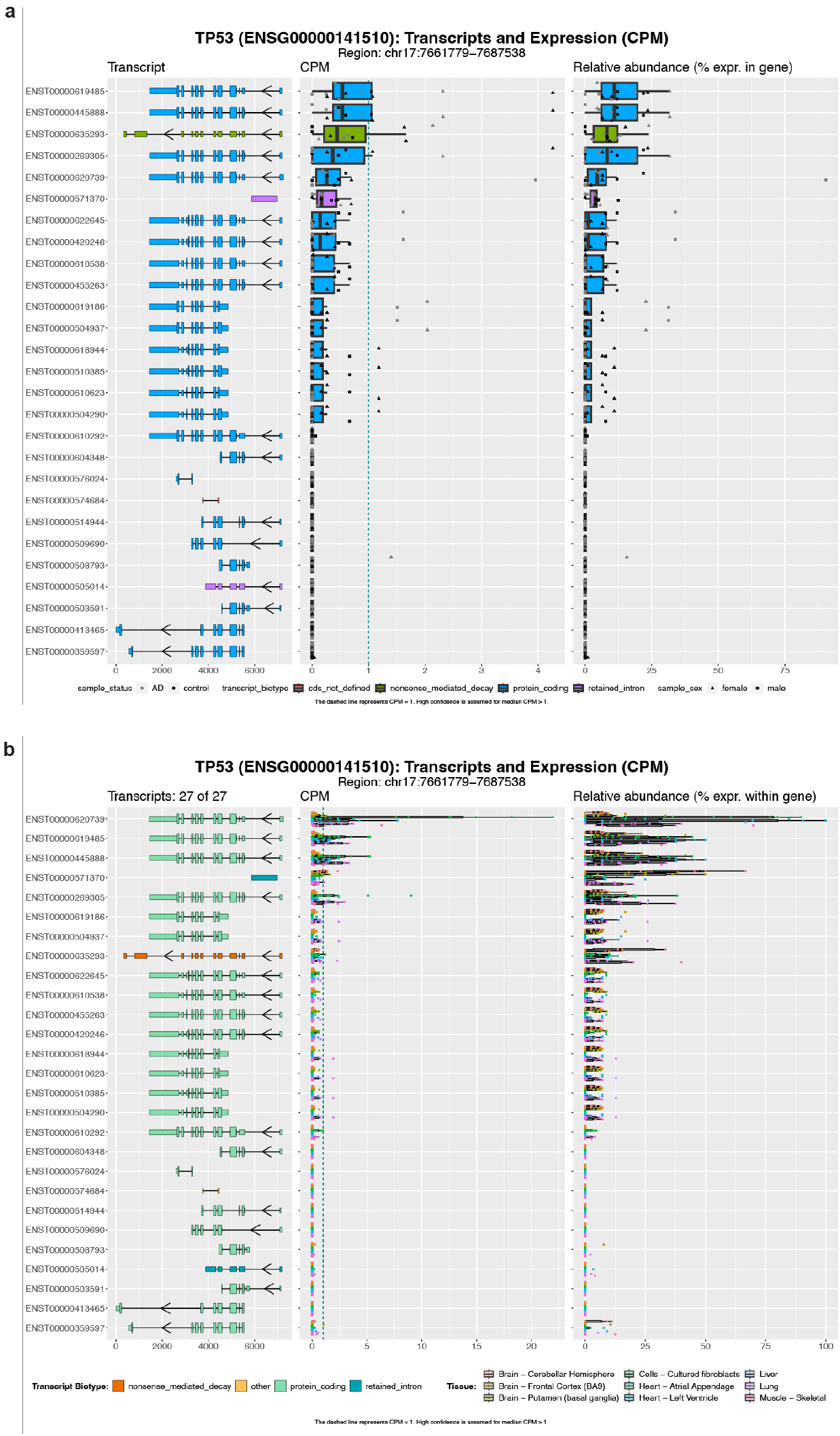

Supplemental Fig. 15. TP53 RNA isoform expression. (**a**) The top *TP53* isoform expressed in human frontal cortex, per Aguzzoli-Heberle et al. was either ENST00000445888 or ENST00000619485. The mRNA sequence between these isoforms is identical, but with distinct annotated coding sequence. The other reported isoforms did not have significant expression levels. (**b**) ENST00000620739 was the most expressed across nine GTEx tissues, per Glinos et al., based on analyses by Page et al. Isoform expression plots based on human frontal cortex data generated by Aguzzoli-Heberle et al. were generated at <https://ebbertlab.com/brain_rna_isoform_seq.html>. Isoform expression plots based on nine GTEx samples were generated at <https://ebbertlab.com/gtex_rna_isoform_seq.html>; these data were generated by Glinos et al. and analyzed by Page et al. All three isoforms are highly similar—ENST00000445888 and ENST00000619485 share identical mRNA sequence but with different annotated coding sequence, whereas ENST00000619485 and ENST00000620739 have identical coding sequences with distinct 5’UTRs.

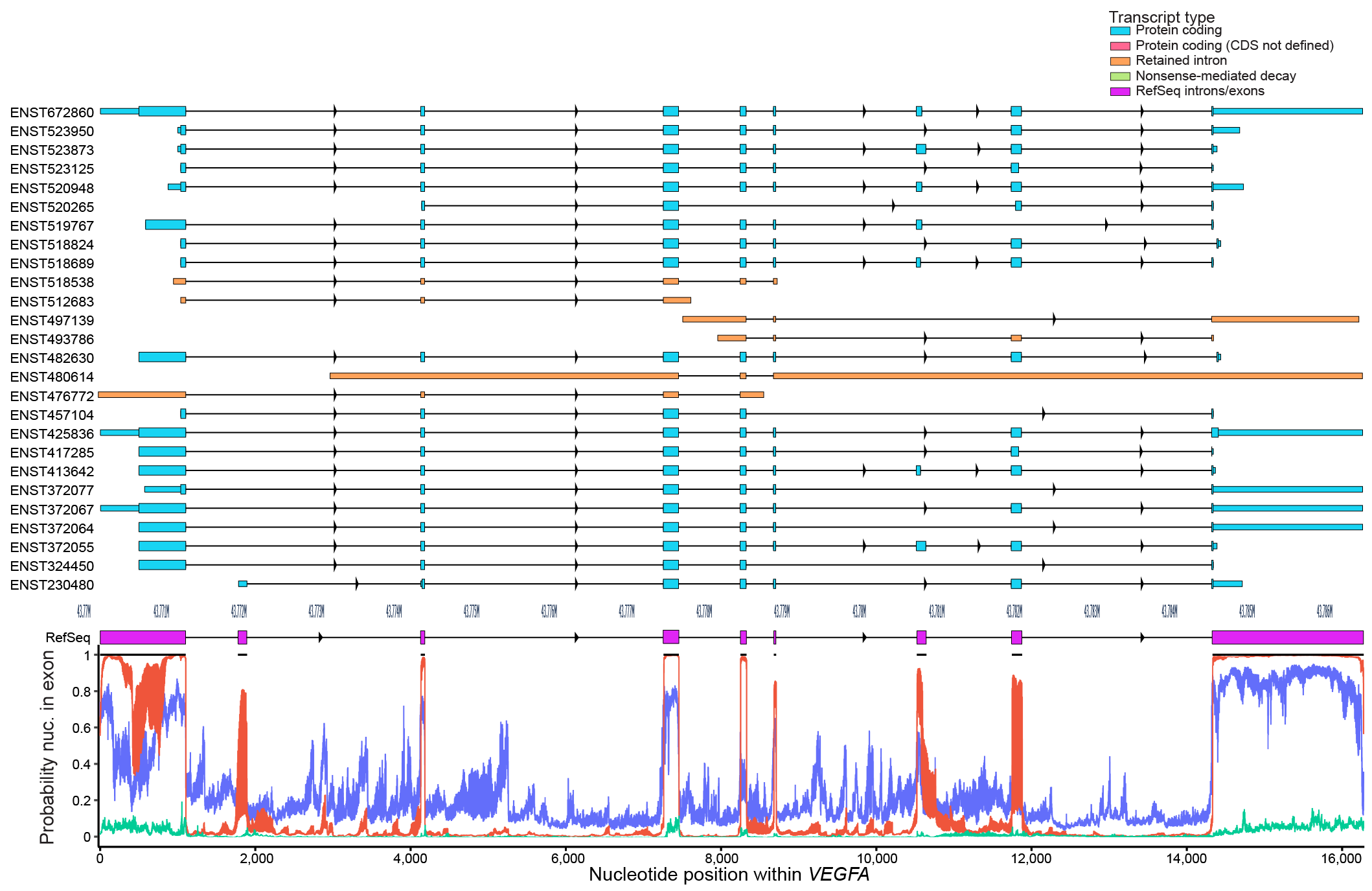

Supplemental Fig. 16. VEGFA RNA isoforms and SegmentNT exonic probabilities. SegmentNT probabilities of being in an exon for *VEGFA*, along with RefSeq exons and all 26 reported Ensembl RNA isoforms.

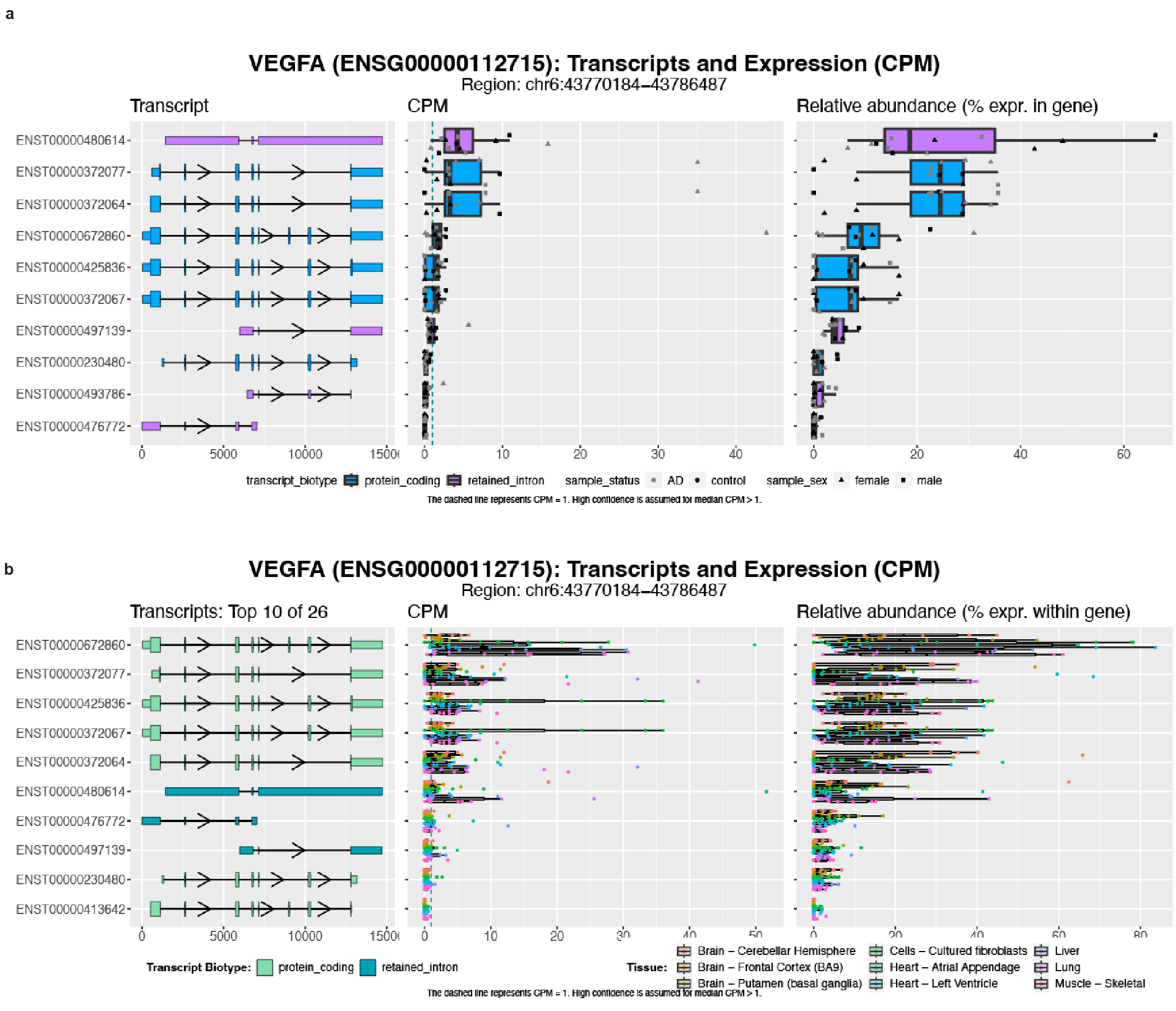

Supplemental Fig. 17. Top ten VEGFA RNA isoform expression. (**a**) The top *VEGFA* isoform expressed in human frontal cortex, per Aguzzoli-Heberle et al. was ENST00000480614, which is reported as an isoform with a retained intron. The other reported isoforms did not have significant expression levels. (**b**) ENST00000672860 was the most expressed across nine GTEx tissues, per Glinos et al., based on analyses by Page et al. Isoform expression plots based on human frontal cortex data generated by Aguzzoli-Heberle et al. were generated at <https://ebbertlab.com/brain_rna_isoform_seq.html>. Isoform expression plots based on nine GTEx samples were generated at <https://ebbertlab.com/gtex_rna_isoform_seq.html>; these data were generated by Glinos et al. and analyzed by Page et al.

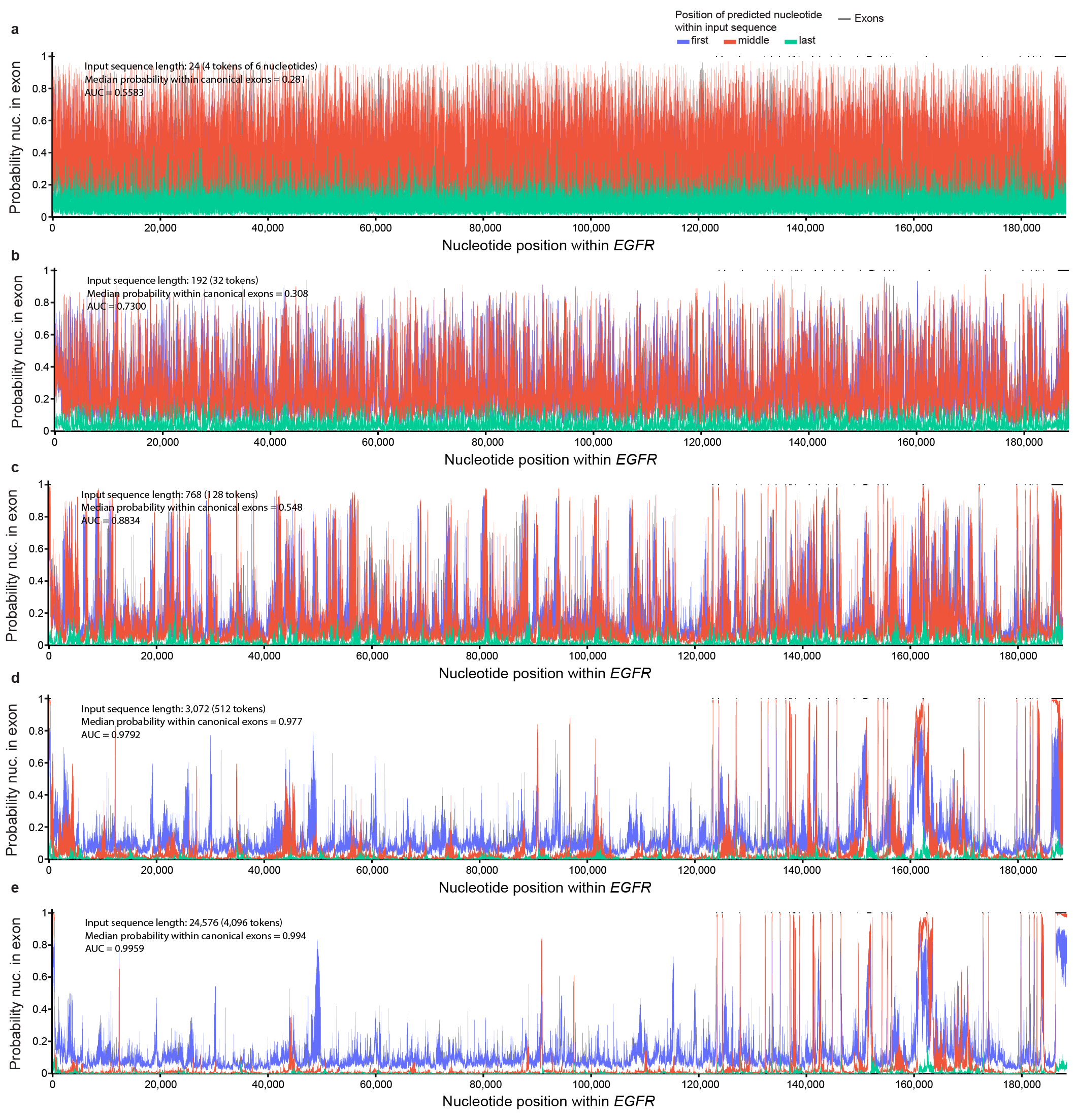

Supplemental Fig. 18. *EGFR* SegmentNT probabilities by input sequence size. (**a**) SegmentNT probabilities for whether each nucleotide is in an exon using an input sequence length of 24 nucleotides (4 tokens). Probabilities appear sporadic with no distinguishable pattern between intronic an exonic nucleotides. (**b**) Same as figure **a**, but input sequences of 192 nucleotides (32 tokens). Probabilities appear less random, but without a clear distinction between intronic and exonic nucleotides. (**c**) Input sequence of 768 nucleotides (128 tokens). Probabilities appear more systematic with a clear pattern distinguishing between intronic and exonic nucleotides. (**d**) Input sequence of 3,072 nucleotides (512 tokens). Probabilities stabilize further. (**e**) Input sequence of 24,576 nucleotides (4,096 tokens). Probabilities for nucleotides within canonical RefSeq exons approach one, while probabilities for nucleotides within canonical RefSeq introns approach zero.

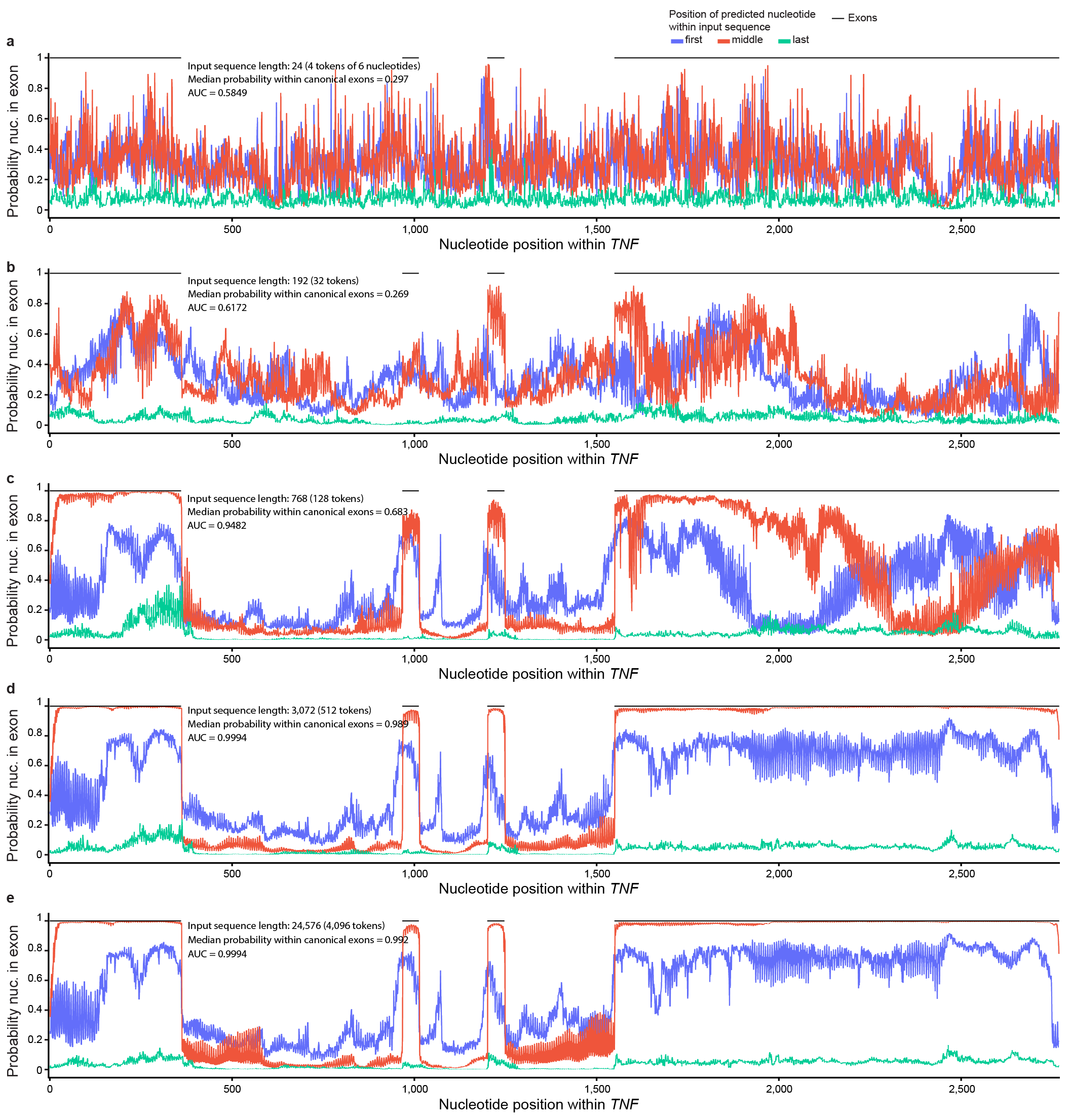

Supplemental Fig. 19. *TNF* SegmentNT probabilities by input sequence size. (**a**) SegmentNT probabilities for whether each nucleotide is in an exon using an input sequence length of 24 nucleotides (4 tokens). Probabilities appear sporadic with no distinguishable pattern between intronic an exonic nucleotides. (**b**) Same as figure **a**, but input sequences of 192 nucleotides (32 tokens). Probabilities appear less random, but without a clear distinction between intronic and exonic nucleotides. (**c**) Input sequence of 768 nucleotides (128 tokens). Probabilities appear more systematic with a clear pattern distinguishing between intronic and exonic nucleotides. (**d**) Input sequence of 3,072 nucleotides (512 tokens). Probabilities stabilize further. (**e**) Input sequence of 24,576 nucleotides (4,096 tokens). Probabilities for nucleotides within canonical RefSeq exons approach one, while probabilities for nucleotides within canonical RefSeq introns approach zero.

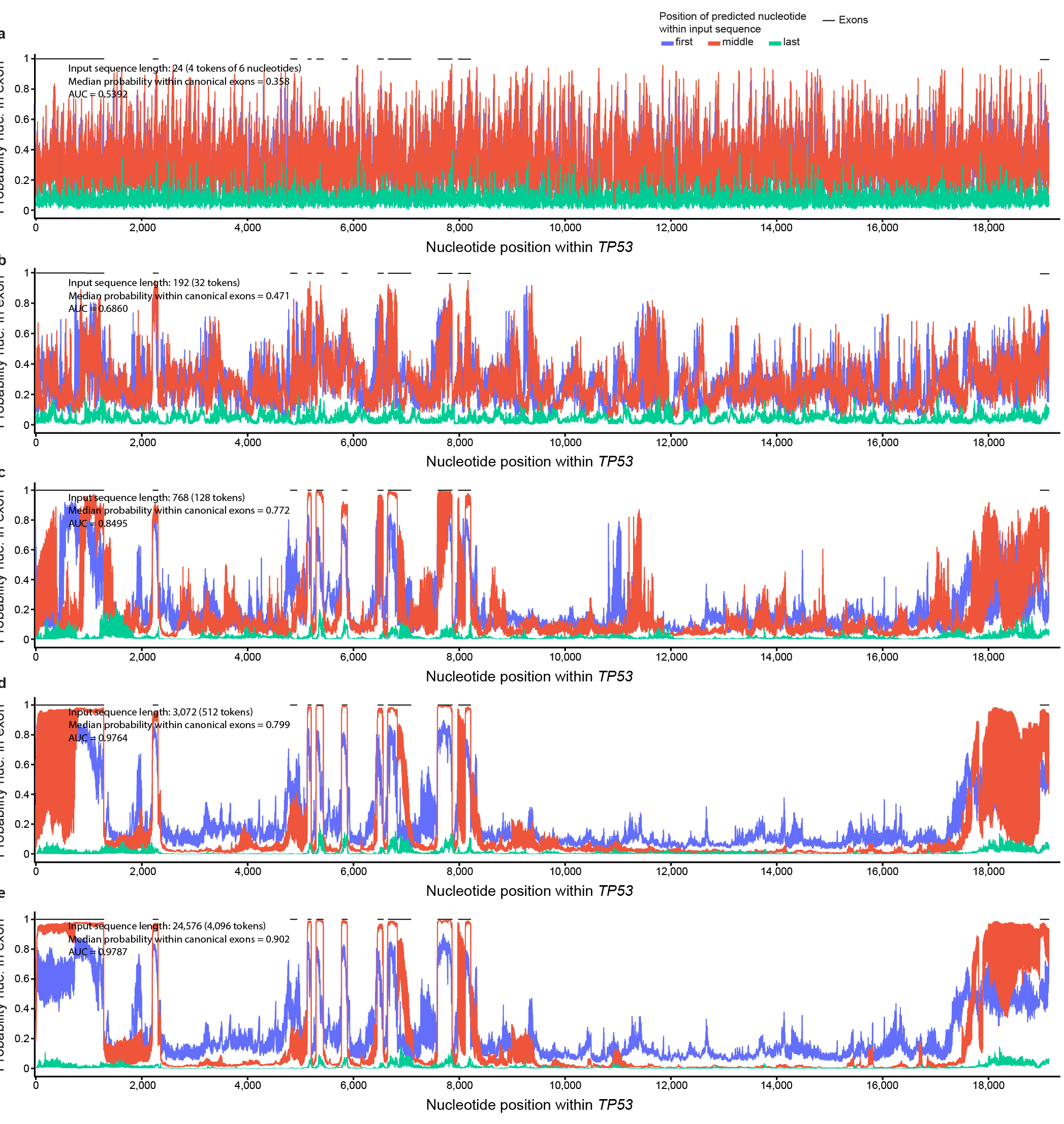

Supplemental Fig. 20. *TP53* SegmentNT probabilities by input sequence size. (**a**) SegmentNT probabilities for whether each nucleotide is in an exon using an input sequence length of 24 nucleotides (4 tokens). Probabilities appear sporadic with no distinguishable pattern between intronic an exonic nucleotides. (**b**) Same as figure **a**, but input sequences of 192 nucleotides (32 tokens). Probabilities appear less random, but without a clear distinction between intronic and exonic nucleotides. (**c**) Input sequence of 768 nucleotides (128 tokens). Probabilities appear more systematic with a clear pattern distinguishing between intronic and exonic nucleotides. (**d**) Input sequence of 3,072 nucleotides (512 tokens). Probabilities stabilize further. (**e**) Input sequence of 24,576 nucleotides (4,096 tokens). Probabilities for nucleotides within canonical RefSeq exons approach one, while probabilities for nucleotides within canonical RefSeq introns approach zero.

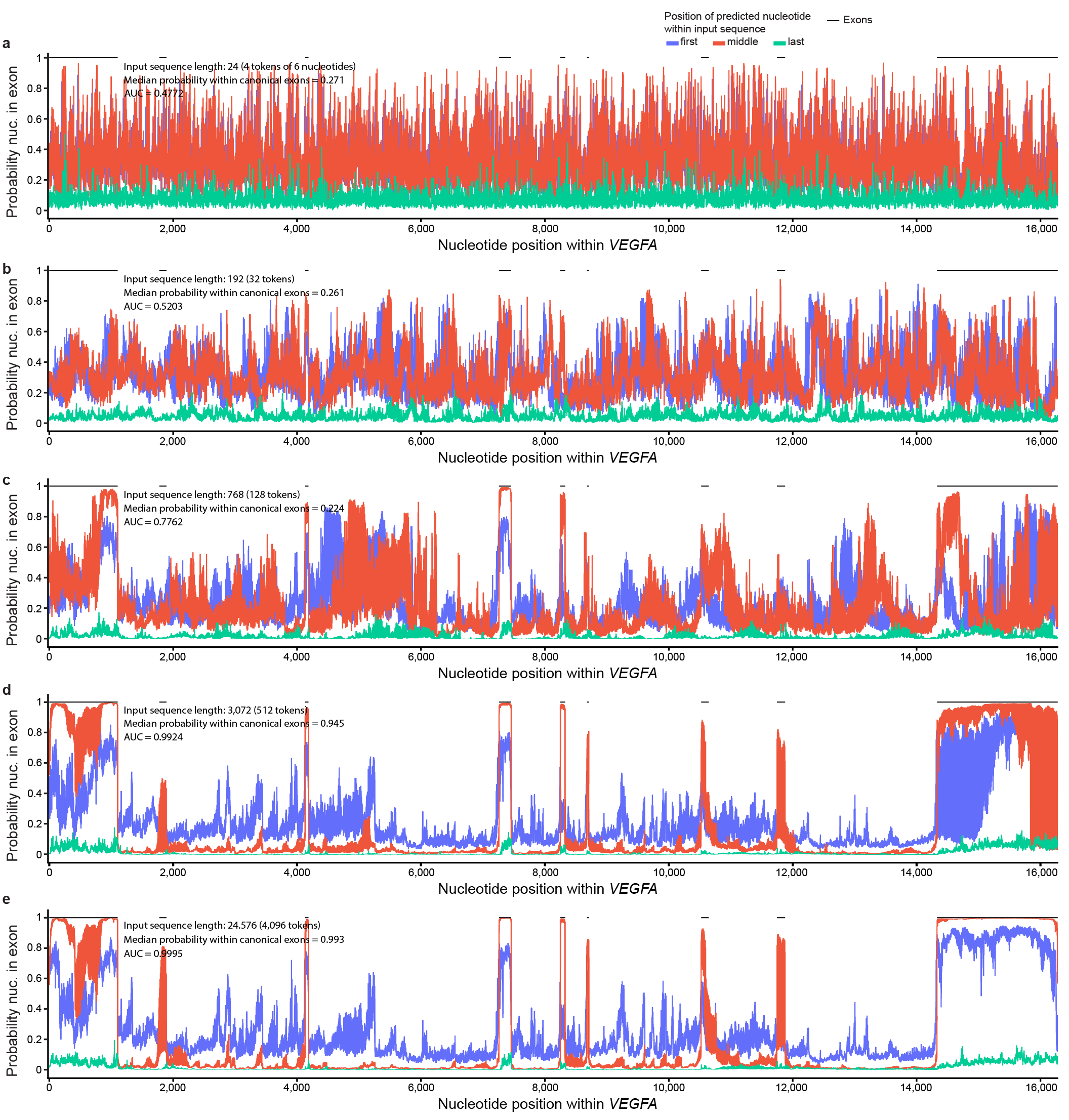

Supplemental Fig. 21. *VEGFA* SegmentNT probabilities by input sequence size. (**a**) SegmentNT probabilities for whether each nucleotide is in an exon using an input sequence length of 24 nucleotides (4 tokens). Probabilities appear sporadic with no distinguishable pattern between intronic an exonic nucleotides. (**b**) Same as figure **a**, but input sequences of 192 nucleotides (32 tokens). Probabilities appear less random, but without a clear distinction between intronic and exonic nucleotides. (**c**) Input sequence of 768 nucleotides (128 tokens). Probabilities appear more systematic with a clear pattern distinguishing between intronic and exonic nucleotides. (**d**) Input sequence of 3,072 nucleotides (512 tokens). Probabilities stabilize further. (**e**) Input sequence of 24,576 nucleotides (4,096 tokens). Probabilities for nucleotides within canonical RefSeq exons approach one, while probabilities for nucleotides within canonical RefSeq introns approach zero.

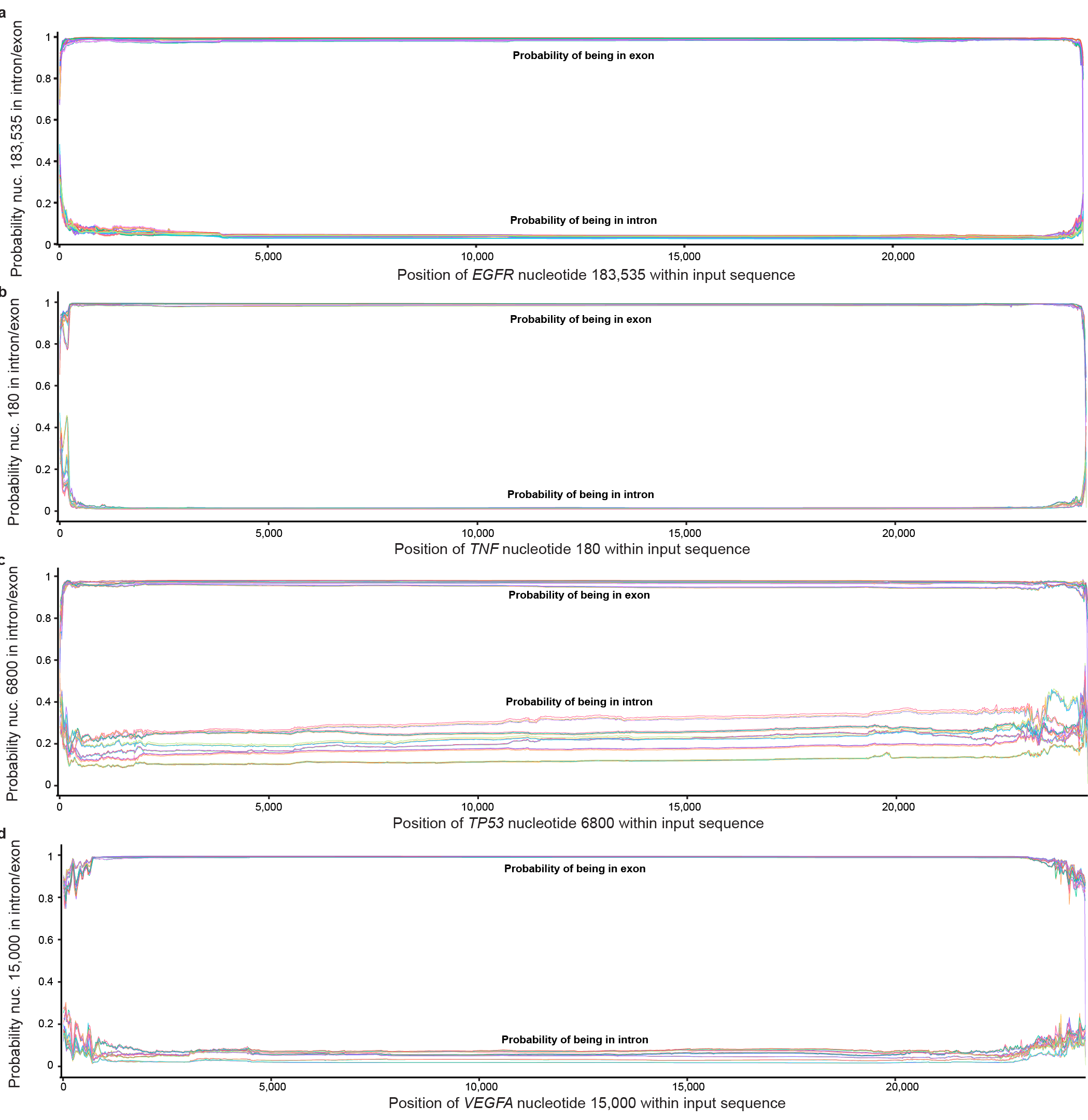

Supplemental Fig. 22. Plotting 24 sets of every 24th position for a randomly selected exonic variant also results in linear, non-cyclical probabilities. As described in main Figure 6, we plotted probabilities in 24 different sets (for both exon and intron predictions), where each set plotted every 24^th^ probability. Specifically, set one plotted probabilities when the selected nucleotide was in positions 0, 24, 48, etc., while set two plotted positions 1, 25, 49, etc. Neither intronic nor exonic probabilities resulted in a cyclical pattern when plotting every 24^th^ value, empirically demonstrating that the cyclical bias operates on a 24-nucleotide cycle.

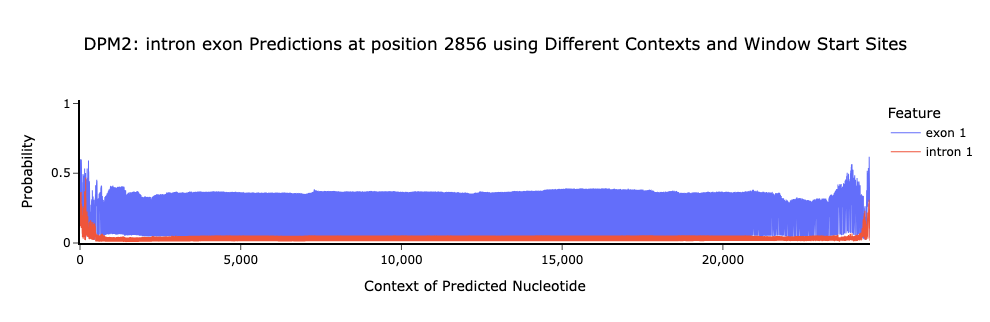

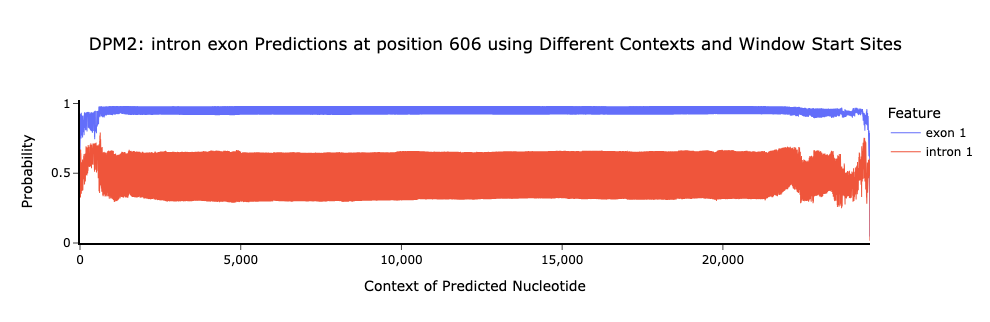

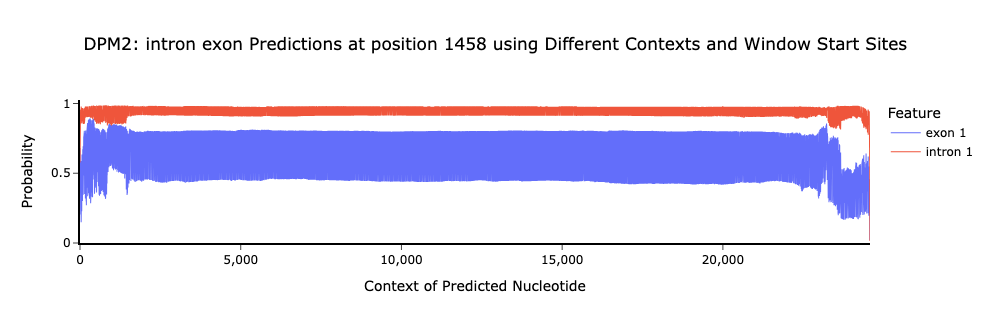

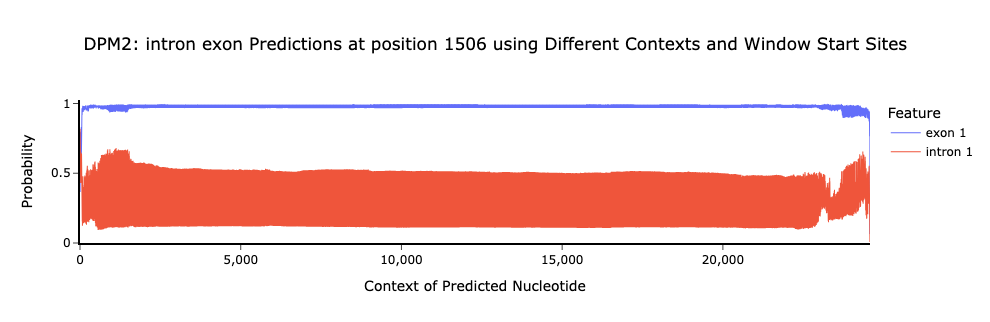

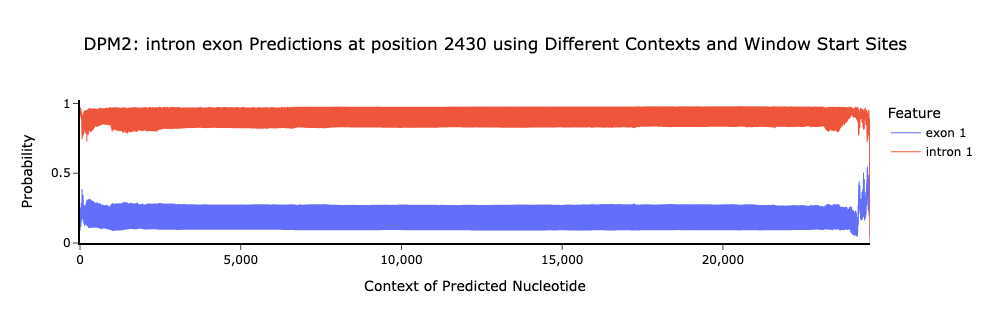

Supplemental Fig. 23. SegmentNT probabilities also oscillate for five *DPM2* nucleotides. As described in main Figure 6, we identified an oscillating pattern for *APOE* nucleotide 850 across every position in the input sequence and even less stable probabilities at distal positions. We observed the same pattern for randomly selected variants in the other four genes in our primary analyses. As further validation, we selected five variants per gene for five additional genes, including *DPM2*, here. In order, we’ve plotted nucleotides 606, 1458, 1506, 2430, and 2856. All exhibit the same oscillating behavior and lower stability at distal positions.

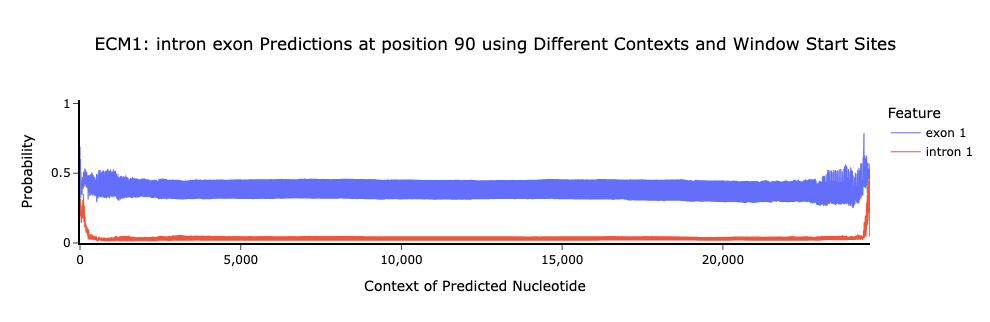

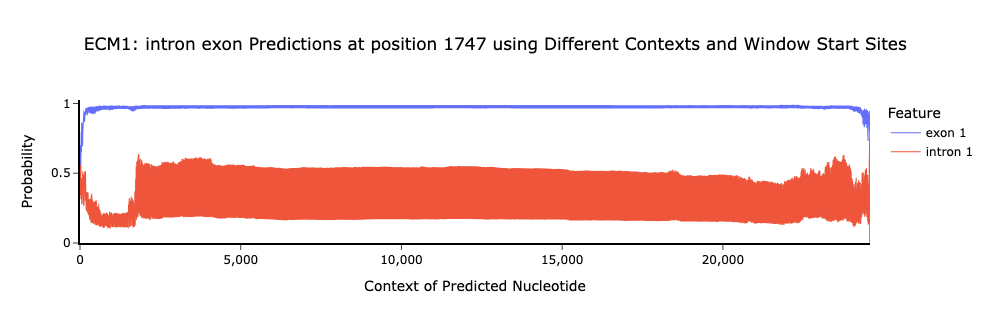

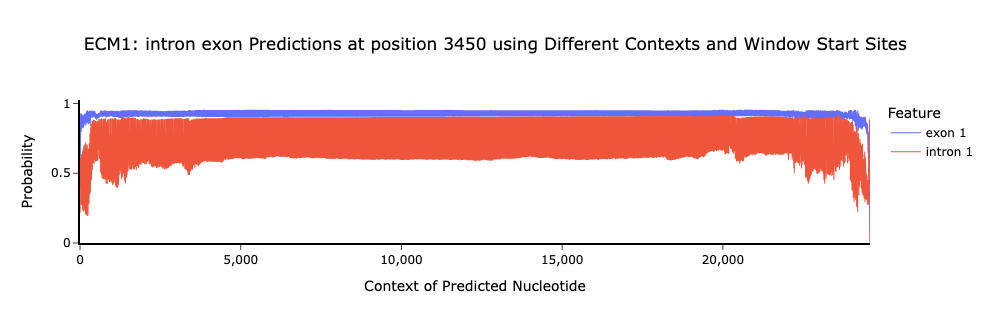

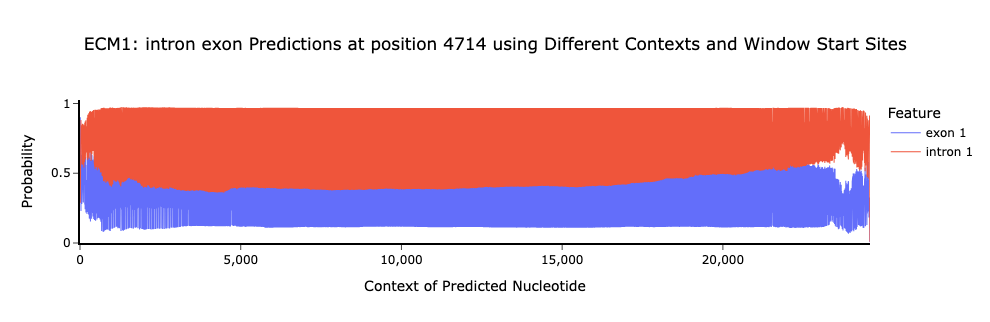

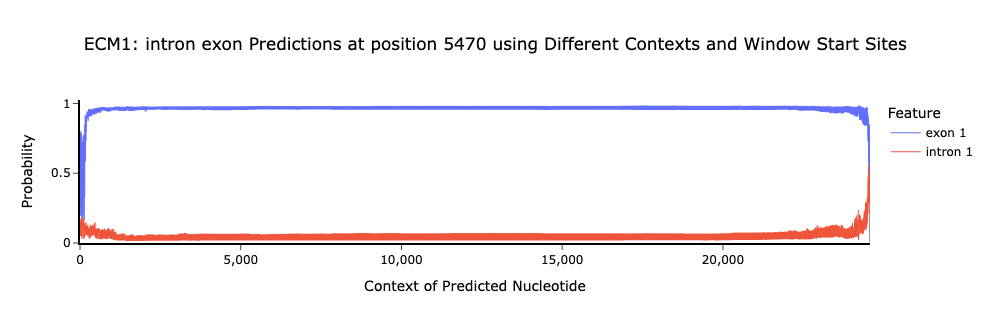

Supplemental Fig. 24. SegmentNT probabilities also oscillate for five *ECM1* nucleotides. As described in main Figure 6, we identified an oscillating pattern for *APOE* nucleotide 850 across every position in the input sequence and even less stable probabilities at distal positions. We observed the same pattern for randomly selected variants in the other four genes in our primary analyses. As further validation, we selected five variants per gene for five additional genes, including *ECM1*, here. In order, we’ve plotted nucleotides 90, 1747, 3450, 4714, and 5470. All exhibit the same oscillating behavior and lower stability at distal positions.

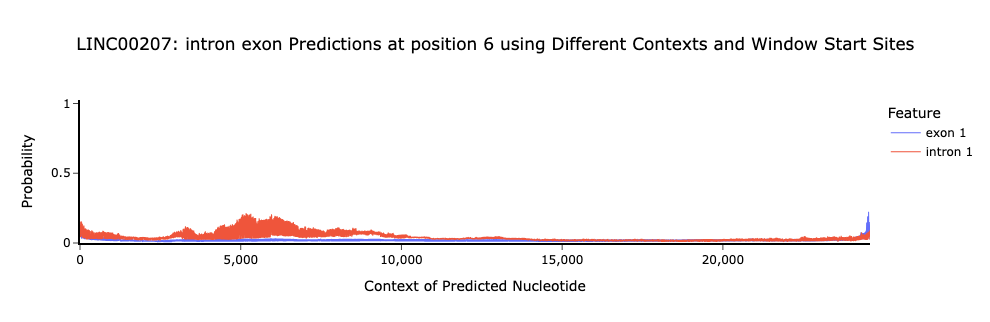

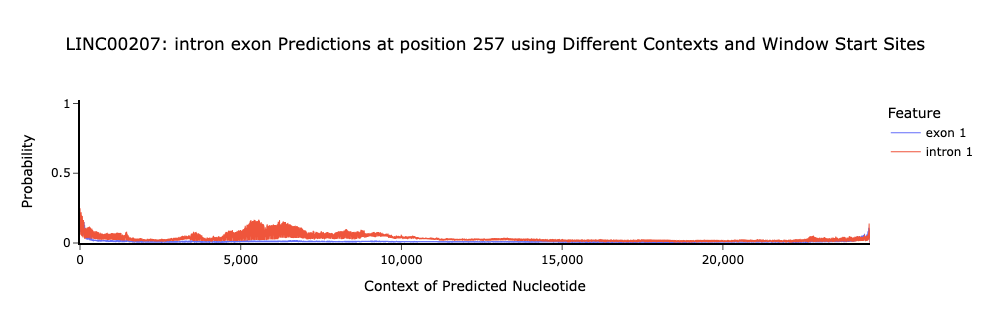

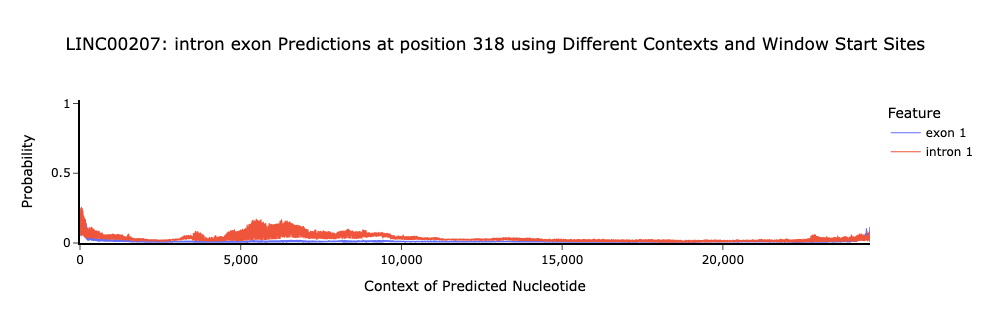

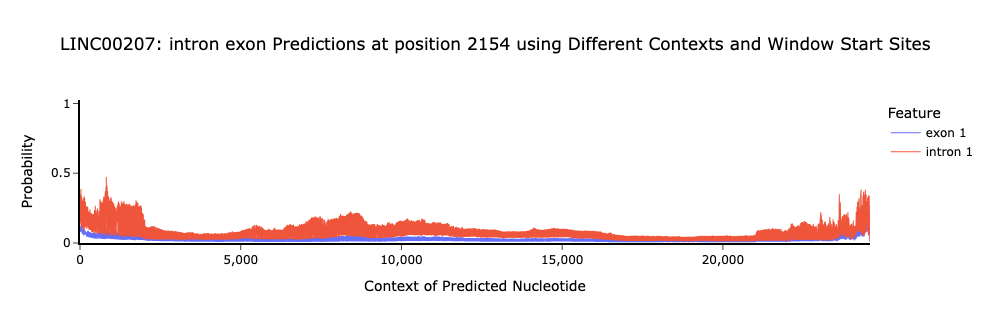

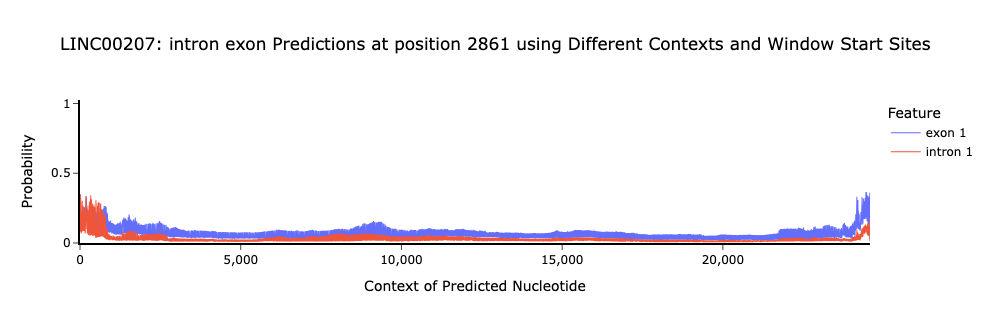

Supplemental Fig. 25. SegmentNT probabilities also oscillate for five *LINC00207* nucleotides. As described in main Figure 6, we identified an oscillating pattern for *APOE* nucleotide 850 across every position in the input sequence and even less stable probabilities at distal positions. We observed the same pattern for randomly selected variants in the other four genes in our primary analyses. As further validation, we selected five variants per gene for five additional genes, including *LINC00207*, here. In order, we’ve plotted nucleotides 6, 257, 318, 2154, and 2861. All exhibit the same oscillating behavior and lower stability at distal positions.

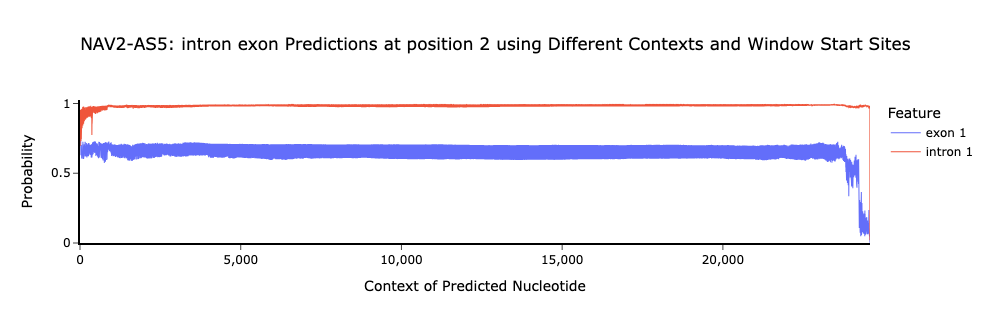

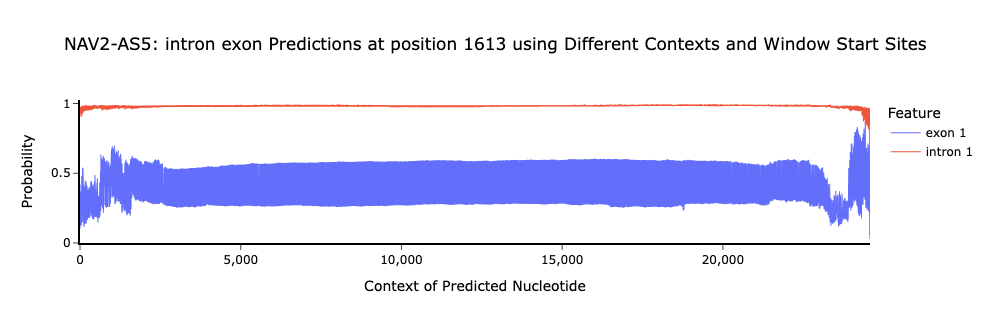

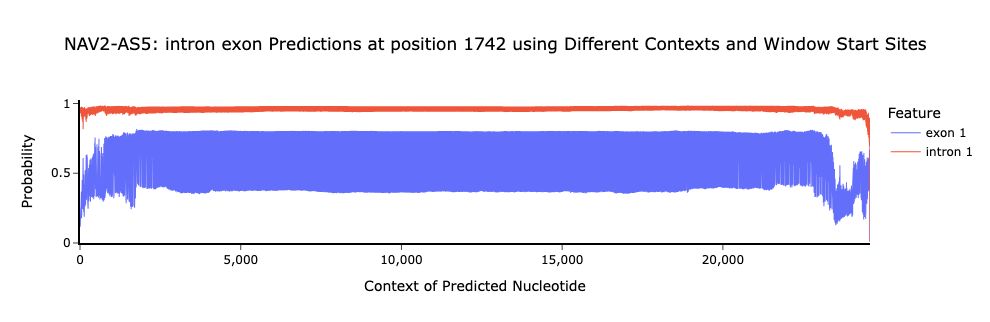

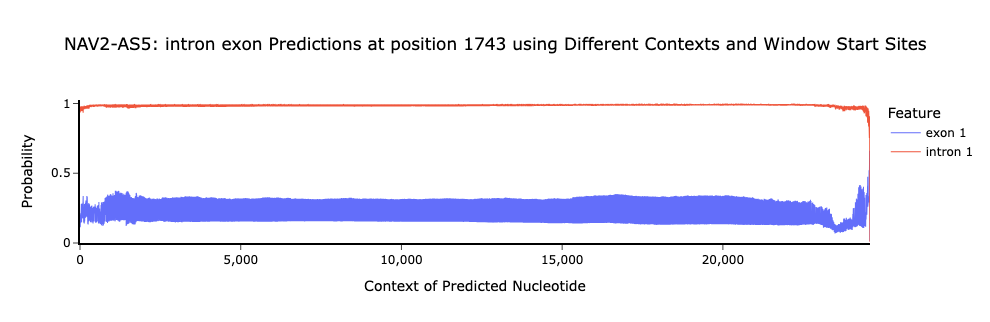

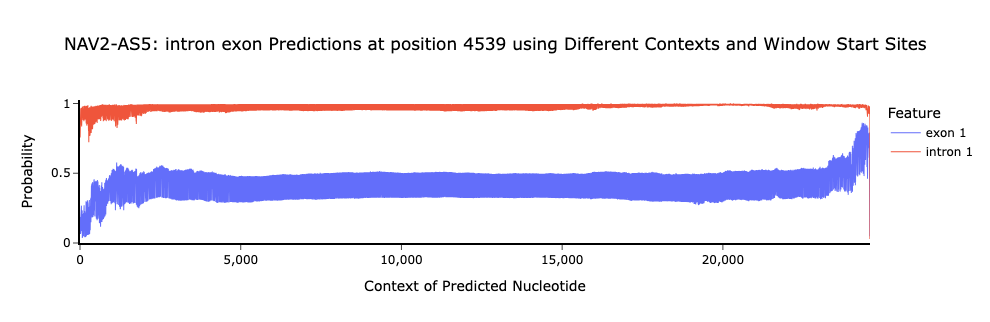

Supplemental Fig. 26. SegmentNT probabilities also oscillate for five *NAV2-AS5* nucleotides. As described in main Figure 6, we identified an oscillating pattern for *APOE* nucleotide 850 across every position in the input sequence and even less stable probabilities at distal positions. We observed the same pattern for randomly selected variants in the other four genes in our primary analyses. As further validation, we selected five variants per gene for five additional genes, including *NAV2-AS5*, here. In order, we’ve plotted nucleotides 2, 1613, 1742, 1743, & 4539. All exhibit the same oscillating behavior and lower stability at distal positions.

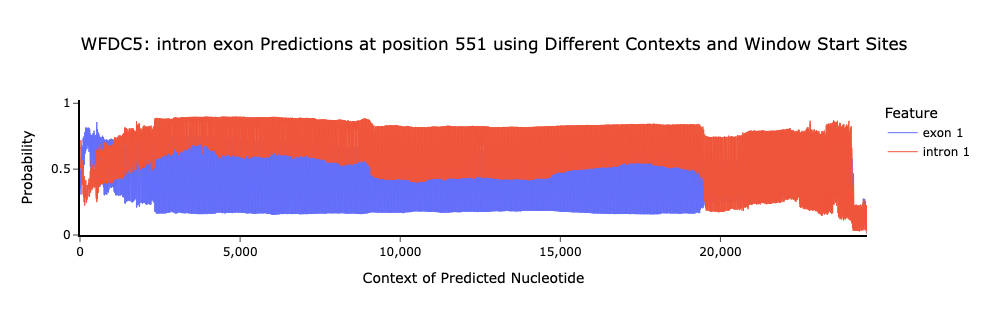

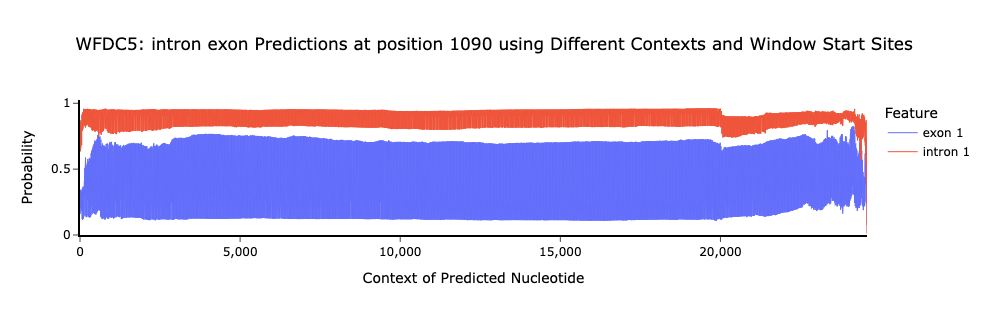

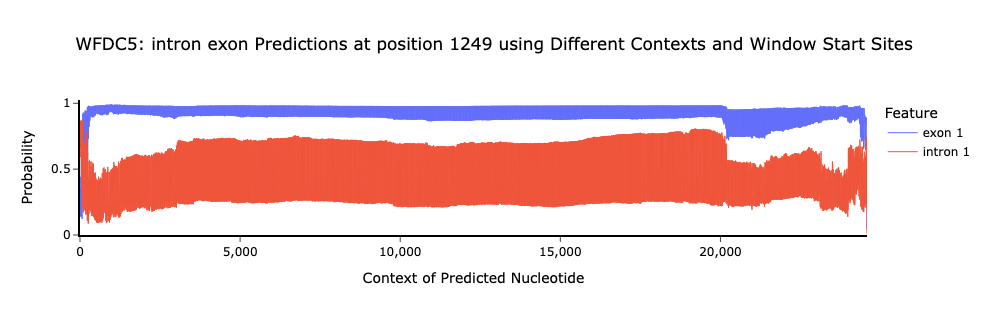

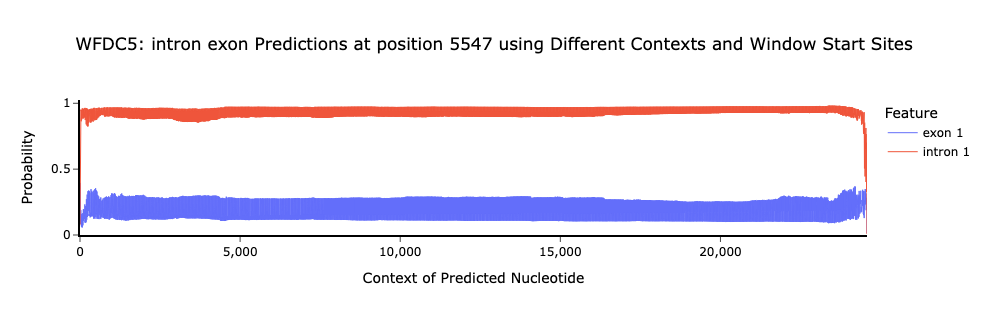

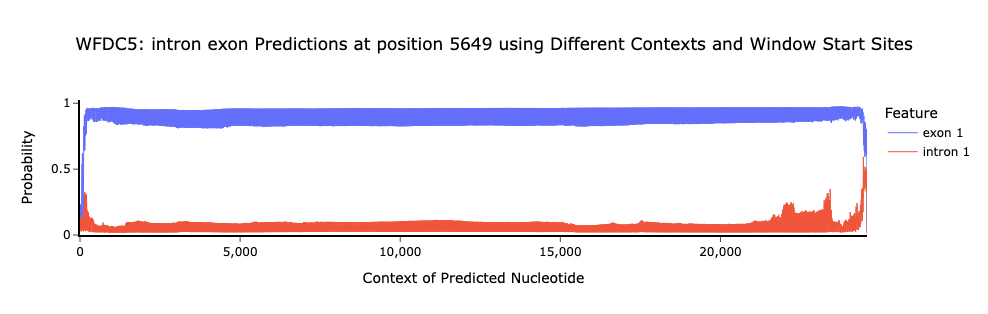

Supplemental Fig. 27. SegmentNT probabilities also oscillate for five *WFDC5* nucleotides. As described in main Figure 6, we identified an oscillating pattern for *APOE* nucleotide 850 across every position in the input sequence and even less stable probabilities at distal positions. We observed the same pattern for randomly selected variants in the other four genes in our primary analyses. As further validation, we selected five variants per gene for five additional genes, including *WFDC5*, here. In order, we’ve plotted nucleotides 551, 1090, 1249, 5547, and 5649. All exhibit the same oscillating behavior and lower stability at distal positions.
