## Supplementary data files for "Systematic contextual biases in SegmentNT potentially relevant to other nucleotide transformer models": generate_isoforms.py

```
import RNApysoforms as RNApy
import plotly.express as px


ensembl_gtf_path = "/Users/mebbert/sequencing_resources/annotations/Homo_sapiens.GRCh38.113.gtf"
annotation = RNApy.read_ensembl_gtf(ensembl_gtf_path)

# APOE #
apoe_annotation = RNApy.gene_filtering(annotation=annotation, target_gene="APOE")
apoe_annotation = RNApy.to_intron(apoe_annotation)
traces = RNApy.make_traces(annotation=apoe_annotation, y='transcript_id',
                           annotation_hue="transcript_biotype",
                           annotation_color_palette=["#18D4F3", "#FFA25A", "#FF6792", "#B7E980"])
fig = RNApy.make_plot(traces = traces, subplot_titles = ["Transcript Structure"], width=1200, height=500)
fig.update_xaxes(showticklabels=True, dtick=100, row=1, col=1)
fig.write_image("./RNApysoforms_APOE_human.svg", format="svg", engine="kaleido", width=5200, height=500)

egfr_annotation = RNApy.gene_filtering(annotation=annotation, target_gene="EGFR")
egfr_annotation = RNApy.to_intron(egfr_annotation)
traces = RNApy.make_traces(annotation=egfr_annotation, y='transcript_id', annotation_hue="transcript_biotype",
                           annotation_color_palette=["#18D4F3", "#FFA25A", "#FF6792", "#B7E980"])
fig = RNApy.make_plot(traces = traces, subplot_titles = ["Transcript Structure"], width=1200, height=500)
fig.update_xaxes(showticklabels=True, dtick=1000, row=1, col=1)
fig.write_image("./RNApysoforms_EGFR_human.svg", format="svg", engine="kaleido", width=5200, height=500)

tnf_annotation = RNApy.gene_filtering(annotation=annotation, target_gene="TNF")
tnf_annotation = RNApy.to_intron(tnf_annotation)
traces = RNApy.make_traces(annotation=tnf_annotation, y='transcript_id', annotation_hue="transcript_biotype",
                           annotation_color_palette=["#18D4F3", "#FFA25A", "#FF6792", "#B7E980"])
fig = RNApy.make_plot(traces = traces, subplot_titles = ["Transcript Structure"], width=1200, height=500)
fig.update_xaxes(showticklabels=True, dtick=1000, row=1, col=1)
fig.write_image("./RNApysoforms_TNF_human.svg", format="svg", engine="kaleido", width=5200, height=500)

tp53_annotation = RNApy.gene_filtering(annotation=annotation, target_gene="TP53")
tp53_annotation = RNApy.to_intron(tp53_annotation)
traces = RNApy.make_traces(annotation=tp53_annotation, y='transcript_id', annotation_hue="transcript_biotype",
                           annotation_color_palette=["#18D4F3", "#FFA25A", "#FF6792", "#B7E980"])
fig = RNApy.make_plot(traces = traces, subplot_titles = ["Transcript Structure"], width=1200, height=500)
fig.update_xaxes(showticklabels=True, dtick=1000, row=1, col=1)
fig.write_image("./RNApysoforms_TP53_human.svg", format="svg", engine="kaleido", width=5200, height=500)

vegfa_annotation = RNApy.gene_filtering(annotation=annotation, target_gene="VEGFA")
vegfa_annotation = RNApy.to_intron(vegfa_annotation)
traces = RNApy.make_traces(annotation=vegfa_annotation, y='transcript_id', annotation_hue="transcript_biotype",
                           annotation_color_palette=["#18D4F3", "#FFA25A", "#FF6792", "#B7E980"])
fig = RNApy.make_plot(traces = traces, subplot_titles = ["Transcript Structure"], width=1200, height=500)
fig.update_xaxes(showticklabels=True, dtick=1000, row=1, col=1)
fig.write_image("./RNApysoforms_VEGFA_human.svg", format="svg", engine="kaleido", width=5200, height=500)
```
