## Supplementary data files for "Systematic contextual biases in SegmentNT potentially relevant to other nucleotide transformer models": findStability.py

```
import ast
import sys
print(sys.argv[1])
#filename = "dpm2_pos1506.txt"   # path to your file
filename=sys.argv[1] #"../../positions/apoe_4096_850.txt"
threshold = 0.8
window_size = 6 #24

with open(filename) as f:
    for line in f:
        line = line.strip()
        if line.startswith("exon="):
            values = ast.literal_eval(line.split("=", 1)[1])
            n = len(values)

            if n < window_size:
                print("List shorter than window size")
                break

            # Compute sliding window averages
            window_avgs = [
                sum(values[i:i + window_size]) / window_size
                for i in range(n - window_size + 1)
            ]

            longest_start = longest_end = None
            longest_len = 0
            current_start = None

            for i, avg in enumerate(window_avgs):
                if avg >= threshold:
                    if current_start is None:
                        current_start = i
                else:
                    if current_start is not None:
                        run_len = i - current_start
                        if run_len > longest_len:
                            longest_len = run_len
                            longest_start = current_start +1
                            longest_end = i  #window_size #since i is already 1 past the end, this encapsulates +1 for 0-base start
                        current_start = None

            # Handle run that reaches the end
            if current_start is not None:
                run_len = len(window_avgs) - current_start
                if run_len > longest_len:
                    longest_start = current_start
                    longest_end = len(window_avgs) - 1

            if longest_start is not None:
                start_index = longest_start
                end_index = longest_end + window_size - 1

                print(f"Longest sliding-window run ≥ {threshold}:")
                print(f"Start index: {start_index}")
                print(f"End index:   {end_index}")
            else:
                print(f"No sliding-window averages ≥ {threshold}")

            break
```
